# Total RNA Sequencing Reveals Early Coding, Noncoding, and Isoform-Level Transcriptomic Changes Following Traumatic and Glaucomatous Optic Nerve Injury

**DOI:** 10.64898/2026.09.08.750121

**Authors:** Noah C. Mathew, Kevin K. Park

## Abstract

**Purpose:** Retinal ganglion cell (RGC) death following optic nerve injury is driven by transcriptional programs involved in apoptosis, neuroinflammation, and other regulated cell death pathways; however, the early transcriptomic changes initiating these processes remain poorly defined. We used rRNA-depleted total RNA sequencing to characterize early alterations in protein-coding genes, long noncoding RNAs (lncRNAs), and alternative splicing events in mouse retinas following optic nerve crush (ONC) and silicone oil-induced ocular hypertension under-detected (SOHU), prior to substantial RGC loss.

**Methods:** Whole retinas were collected 6 hours after ONC and 5 days after SOHU induction. Differentially expressed protein-coding genes and lncRNA-classified transcripts were identified using total RNA sequencing, followed by functional enrichment, lncRNA-associated pathway analysis, alternative splicing analysis, and fluorescent in situ hybridization (FISH) validation.

**Results:** In SOHU retinas, hundreds of protein-coding genes and lncRNA-classified transcripts were differentially expressed, with enrichment of immune activation, extracellular matrix remodeling, complement, phagocytic signaling, and suppression of mitochondrial energy metabolism pathways. Similarly, in ONC retinas, many protein-coding genes and lncRNA-classified transcripts were differentially expressed, with a prominent chemokine, cytokine-dominant inflammatory response. FISH validation supported injury-associated upregulation of *Prss56* and *Lif*. Both injury models exhibited substantial transcript-level alterations, with widespread alternative splicing changes detected.

**Conclusions:** Early optic nerve injury induces rapid transcriptional changes involving protein-coding genes, lncRNAs, and transcript isoform regulation before substantial RGC loss. These findings highlight early alterations in both coding and noncoding RNA landscapes following retinal injury and provide a valuable resource of candidate genes and pathways for future validation in RGC degeneration and neuroprotection.

## INTRODUCTION

Optic nerve injury is a major cause of irreversible visual impairment and serves as a widely used model for studying neurodegeneration within the central nervous system (CNS).^1–4^ Retinal ganglion cells (RGCs), whose axons form the optic nerve and relay visual information from the retina to the brain, are highly vulnerable to axonal damage.^5^ Following traumatic injury, or glaucomatous insult, injured axons undergo degeneration, and a substantial proportion of RGCs subsequently die, resulting in permanent loss of visual function due to the limited regenerative capacity of adult CNS neurons.^2,5^

RGC death after axonal injury is mediated by multiple molecular pathways. Apoptosis, particularly through the intrinsic BAX-dependent and caspase-mediated pathway, has long been considered a central mechanism of neuronal loss following optic nerve injury.^6,7^ However, accumulating evidence indicates that additional forms of regulated cell death, including necroptosis, pyroptosis, and ferroptosis, also contribute to RGC degeneration.^8–10^ Furthermore, neuroinflammation, mitochondrial dysfunction, oxidative stress, and loss of target-derived trophic support may interact with these death pathways to determine neuronal fate.^11,12^ Understanding the mechanisms that link axonal injury to RGC death has significant clinical relevance for diseases such as glaucoma, traumatic optic neuropathy, ischemic optic neuropathy, and multiple sclerosis-associated optic neuritis.^2,5,13,14^ Despite considerable progress in identifying downstream cell death pathways, the earliest molecular events that trigger the degenerative response after axonal injury remain incompletely understood. Defining these initial signaling mechanisms is critical for developing effective neuroprotective therapies capable of preventing RGC loss before irreversible degeneration occurs.

Previous transcriptomic studies of mouse retinal injury models have identified changes in genes associated with inflammation, stress responses, axonal damage, neuronal survival, and regenerative potential.^15–17^ However, much of this work has focused primarily on protein-coding genes, while the contribution of noncoding transcripts, including long noncoding RNAs (lncRNAs), remains less well understood. In addition, many transcriptomic studies have focused on later stages after injury, when RGC degeneration and cell death pathways are already activated. Consequently, the earliest transcriptional responses that initiate neuronal degeneration remain poorly defined.

LncRNAs are increasingly recognized as critical regulators of diverse cellular processes, including gene transcription, chromatin remodeling, RNA splicing and stability, epigenetic regulation, and cellular stress responses.^18,19^ Through interactions with DNA, RNA, and protein complexes, lncRNAs can influence gene expression programs at multiple levels and contribute to both physiological homeostasis and disease pathogenesis.^18–20^ In the CNS, numerous lncRNAs have been implicated in neuroinflammation, and neurodegeneration.^21^ Emerging evidence suggests that similar mechanisms operate within the retina, where lncRNAs may regulate neuronal survival, glial activation, and responses to injury.^10,18,19,22^ Recent studies have potentially linked dysregulated lncRNA expression to RGC apoptosis, axonal degeneration, oxidative stress, and inflammatory signaling in models of optic neuropathy and other neurodegenerative retinal disorders.^10,18,19,22,23^ These findings point to lncRNAs as potentially important mediators of the molecular events that drive retinal damage and repair following optic nerve injury.

Despite growing interest in the role of lncRNAs in retinal disease, the temporal dynamics and functional significance of lncRNA expression during the early phase of optic nerve injury remain poorly understood. Most studies have focused on later stages of degeneration or on individual candidate lncRNAs, leaving the broader transcriptomic landscape largely unexplored during the critical initial period when injury-induced signaling pathways are activated and cellular fate decisions are established. A comprehensive characterization of early lncRNA responses may therefore provide important insights into the molecular mechanisms underlying RGC injury, identify novel regulators of neurodegenerative pathways, and reveal potential therapeutic targets for preserving vision after optic nerve damage.

There is growing interest in the application of single-cell RNA sequencing (scRNA-seq) and spatial transcriptomics, which provide valuable cell type-specific and spatially resolved insights into the CNS, retinal biology, and disease.^24,25^ However, many widely used scRNA-seq methods currently rely on oligo(dT)-based capture of poly(A) transcripts, limiting detection of many non-polyadenylated RNAs, while the sparse nature of single-cell datasets can reduce sensitivity for low-abundance transcripts, including numerous lncRNAs.^26,27^

To obtain a comprehensive view of the earliest molecular responses to optic nerve injury, we performed ribosomal RNA (rRNA)-depleted total RNA sequencing (hereafter referred to as total RNA-seq) of mouse retinas at very early time points following optic nerve crush (ONC) (i.e., 6 hours post crush) and in the silicone oil induced hypertension under-detected (SOHU) glaucoma model (i.e., 5 days post SOHU induction), prior to substantial RGC loss.^28–31^ Unlike conventional poly(A)-selected RNA-seq, which primarily captures polyadenylated mRNAs, total RNA-seq enables simultaneous profiling of both protein-coding transcripts and a broad spectrum of noncoding RNAs, including polyadenylated and non-polyadenylated lncRNAs.^32,33^ This approach provides a more complete assessment of injury-induced transcriptional changes and is particularly well suited for identifying previously unrecognized noncoding regulators.^32,33^

Through this work, we identified differentially expressed protein-coding genes and lncRNA-classified transcripts. We performed functional enrichment analyses, examined putative lncRNA-associated pathways, and assessed differential alternative splicing using RNA-seq data. Candidate transcript changes were further evaluated using fluorescence in situ hybridization (FISH). Together, this study provides an early whole-retina transcriptomic comparison of ONC and SOHU injury models and highlights noncoding RNA dysregulation and isoform-level transcript remodeling as potential regulatory features of retinal injury. Notably, robust transcriptomic changes were detectable as early as 6 hours after optic nerve crush, underscoring the remarkably rapid molecular response of the retina to axonal injury and suggesting that critical regulatory programs are initiated within hours of injury onset.

## MATERIALS AND METHODS

### Experimental methods statement

All experimental methods were in full compliance with the most recent regulations and guidelines provided by the University of Texas Southwestern (UTSW) Medical Center.

### Animals

Experimental procedures conducted were compliant with all animal use protocols provided by The UTSW Medical Center’s Institutional Animal Care and Use Committee (IACUC) and adhered to the ARVO Statement for the Use of Animals in Ophthalmic and Vision Research. Eight-week-old C57BL/6J WT male mice (Strain #000664) were purchased from Jackson Laboratories (Bar Harbor, Maine) and utilized for both ONC and SOHU experiments. Experimental and control mice were generated at the same relative time to ensure age matched replicates and experimental groups. Only male mice were used to minimize transcriptional variability associated with hormonal fluctuations across the estrous cycle ^34,35^ thereby improving the detection of subtle, injury-induced transcriptional changes.

### ONC procedure

ONC was conducted as described in previous publications.^36^ Specifically, eight-week-old C57BL/6J male mice were anesthetized via isoflurane (3% isoflurane at 2L/min). An incision was made on the superior posterior position of the conjunctiva of the oculus sinister (OS) where the optic nerve was found behind the globe after carefully moving aside muscle and tissue. Crush was conducted using a fine Dumont # 5 forceps (Fisher Scientific, Hampton, New Hampshire) for 15 seconds. For sham procedure, the optic nerve was exposed but not crushed. Oculus dexter (OD) of both experimental and sham procedures had no surgery conducted. Intraperitoneal injections of buprenorphine (0.3 mg/ml) were administered to mice as an analgesic post procedure. Artificial tears were administered to mice to prevent drying of the eyes or cataract formation. Mice with excessive bleeding were excluded from downstream analysis.

### SOHU procedure

SOHU procedure was conducted as described in previous publications.^28,37^ Eight-week-old C57BL/6J male mice were anesthetized through an intraperitoneal injection of Avertin (0.3 mg/g). A 32-gauge lancet (Owen Mumford, Chipping Norton, United Kingdom) was used to initially puncture through the layers of the cornea allowing for the aqueous humor to drain out of the anterior chamber for 1 min through the tunneling site. After draining of the aqueous humor, 2 µl of silicone oil (Alcon Silicon Oil 1000, Medex Supply, Passaic, NJ, USA) was injected into the anterior chamber using a 34G needle (JBP Ultra-Thin Wall Nanoneedle, 34G × 4 mm, AD Surgical, Sunnyvale, CA, USA). Size of the oil droplet must be > 1.8mm for mice to be used for further analysis. Mice with damaged iris or pupil were not used for downstream analysis.

Injection of ∼2 µl of PBS (saline solution) was used for sham procedure. All experimental procedures were conducted in the OS, where OD had no experimental procedures conducted. Intraperitoneal injections of buprenorphine (0.3 mg/ml) were administered to mice as an analgesic post procedure. Artificial tears were administered to mice to prevent drying of the eyes or cataract formation.

### Intraocular pressure (IOP) checks

IOP was measured using a rebound tonometer following the protocol described in the SOHU model developed by.^28,37^ Mice were anesthetized via isoflurane (3% isoflurane at 2L/min). Because the SOHU model produces ocular hypertension primarily in the posterior and vitreous chambers due to silicone oil (SO)–induced pupillary block, IOP measurements were standardized at approximately 15 minutes after induction of anesthesia, when pupil dilation reconnects the anterior and posterior chambers and allows accurate detection of pressure changes with rebound tonometry.^28^ IOP measurements were obtained using a tonometer (TonoLab, Colonial Medical Supply, Espoo, Finland) according to the manufacturer’s instructions. Each automated measurement consisted of 6 probe contacts with the central cornea; the device discarded the highest and lowest values and calculated the mean of the remaining readings to generate a single machine-derived value. Ten independent machine-generated values were obtained for each eye, where average of these values was used as the final IOP measurement. Baseline IOP was measured prior to SOHU/SHAM induction, and post-induction IOP was measured 5 days after induction.

### Retinal RNA extraction and quality control

Total RNA was isolated from whole mouse retinas using the Qiagen RNeasy Mini Kit (Qiagen, Hilden, Germany) according to the manufacturer’s protocol. Mice were euthanized and eyes were rapidly enucleated. Under a dissecting microscope, retinas were carefully dissected from the eyecups and immediately placed into RNase-free microcentrifuge tubes on ice containing 350 µL of RLT buffer supplemented with β-mercaptoethanol (1%). Retinal tissues were mechanically homogenized using a bead-mill homogenizer (TissueLyser II, Qiagen, Hilden, Germany). The homogenate was briefly centrifuged to remove debris and mixed with 350 µL of 70% ethanol to facilitate RNA binding to the silica membrane. The lysate was then transferred to a RNeasy spin column and centrifuged at 8,000 × g for 15 s to allow RNA binding. Columns were washed sequentially with RW1 buffer followed by two washes with RPE buffer to remove contaminants including proteins and genomic DNA. After the final wash, the column was centrifuged for 1 min at full speed to eliminate residual wash buffer. RNA was eluted in 40 µL of RNase-free water by centrifugation at 8,000 × g for 1 min. RNA concentration and purity were assessed by spectrophotometry using a NanoDrop 2000 Spectrophotometer (Thermo Fisher Scientific, Waltham, MA, USA). RNA integrity and concentration were further evaluated through Qubit and Tapestation analysis. Samples with a RIN score of >7 as well as appropriate concentration of RNA were stored at −80 °C until all samples were ready to be shipped to Novogene for RNA sequencing analysis (Figure 1).

**Figure 1.**
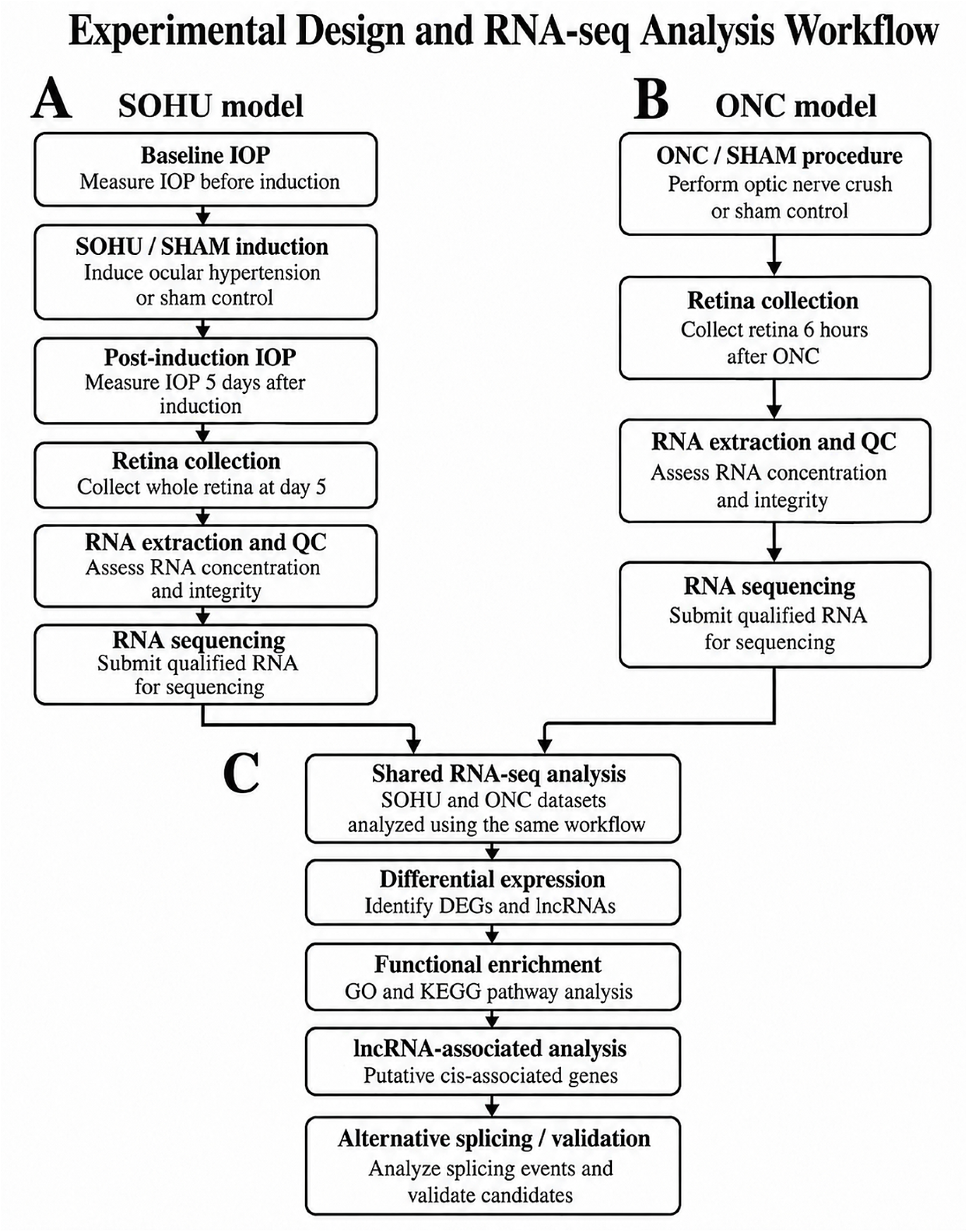
Experimental design and RNA-seq analysis workflow. (**A**) SOHU retinas were collected 5 days after induction following baseline and post-induction IOP measurements. (**B**) ONC retinas were collected 6 hours after optic nerve crush. Retinal RNA was extracted, assessed for quality and concentration using Qubit and TapeStation, and submitted for rRNA depleted total RNA sequencing. (**C**) Sequencing data were analyzed for differential gene and lncRNA expression, functional enrichment, identification of putative cis-associated genes for differentially expressed lncRNAs, alternative splicing, and candidate validation.

### rRNA-depleted total RNA-seq and transcriptomic analysis

RNA sequencing was performed to identify differentially expressed genes (DEGs) and long non-coding RNAs (lncRNAs) in mouse retinal tissue. Total RNA extracted from whole retinas was submitted to Novogene (Sacramento, CA, USA) for library preparation, sequencing, and initial bioinformatic processing using the Novogene Ultra-Low Input Total RNA Sequencing Service (Figure 1). RNA quantity and integrity were re-assessed prior to library preparation to ensure suitability for sequencing. Total RNA was used for sequencing library preparation. Libraries were constructed from low-input total RNA samples using a strand-specific protocol optimized for transcriptome profiling from limited starting material. Ribosomal RNA was removed to enrich for coding and non-coding transcripts, enabling detection of both messenger RNAs (mRNAs) and lncRNAs. The remaining RNA was fragmented under elevated temperature using divalent cations. First-strand cDNA synthesis was performed using random hexamer primers and M-MuLV reverse transcriptase, followed by second-strand cDNA synthesis using DNA Polymerase I and RNase H. cDNA fragments were end-repaired, adenylated at the 3′ ends, ligated to indexed adapters, and size-selected to enrich for fragments approximately 370–420 bp using AMPure XP beads. Libraries were treated with USER Enzyme, PCR-amplified using Phusion High-Fidelity DNA polymerase, purified with AMPure XP beads, assessed for quality using the Agilent 5400 system, and quantified by qPCR. Prepared libraries were quantified, pooled, and sequenced on an Illumina platform (Illumina, San Diego, CA, USA) to generate paired-end reads. Low-quality reads were removed from downstream analysis through filtering of raw read results. Raw reads were filtered by removing reads containing adapters, reads containing N > 0.1% undetermined bases, and low-quality reads in which more than 50% of bases had a quality score (Qscore) of ≤ 5 (Supplementary Figure 1). Sequencing quality was further evaluated using Q20, Q30, overall error rate, and GC-content metrics (Supplementary Figure 2). Clean reads were aligned to the mouse reference genome (GRCm39/mm39) using the splice-aware aligner HISAT2.^38^ Genome-level alignment was performed to support transcript assembly and alternative-splicing analysis, including the detection of splice-junction reads. The resulting alignments were used for downstream genomic-region characterization and transcript assembly. RNA sequencing generated an average read depth of approximately 124 million total reads per sample for the SOHU dataset and 127 million total reads per sample for the ONC dataset. Transcript assembly and quantification were performed to determine gene and transcript abundance. Transcript assembly was performed using StringTie.^39^ Mapping information from all samples was combined and used for transcript assembly and assembled transfrags were compared against reference transcripts to identify novel genes, novel exons of known genes, and refined transcript start/end sites, and mapped RNA-seq reads were classified according to genomic region. (Supplementary Figure 3).

Novel transcript outputs were classified as lncRNA or mRNA based on transcript annotation and structural features, including gene biotype, exon number, transcript length, and open reading frame length. Coding potential was evaluated independently using CPC, CNCI, and PFAM database analyses. Only transcripts classified as noncoding by all three approaches, corresponding to the intersection of the three prediction outputs, were retained as candidate lncRNAs (Supplementary Figure 4A). Feature-signature analyses comparing exon number, transcript length, and ORF length between lncRNAs and mRNAs were used to support classification of noncoding and protein-coding transcripts (Supplementary Figure 4B-4D). Expression levels were calculated as fragments per kilobase of transcript per million mapped reads (FPKM) (Supplementary Figure 5). Read counts were normalized using scaling factors implemented in edgeR, and differential expression between injury and sham groups was analyzed using the edgeR R package.^40^ P values were adjusted for multiple testing using the Benjamini–Hochberg method. Significantly differentially expressed genes and transcripts were defined as those with absolute log2 fold change ≥ 1 and adjusted p-value ≤ 0.05. Gene Ontology (GO) and Kyoto Encyclopedia of Genes and Genomes (KEGG) pathway enrichment analyses were performed on differentially expressed genes identified from both SOHU retinas 5 days after silicone oil injection and ONC retinas 6 hours post crush. Upregulated and downregulated genes were analyzed separately to identify biological processes, molecular functions, cellular components, and pathways associated with SOHU-induced transcriptional changes. GO enrichment included biological process, cellular component, and molecular function categories, while KEGG enrichment was used to identify significantly enriched biological pathways. Enrichment results included the term description, GeneRatio, background ratio, nominal P value, adjusted P value, associated gene IDs/gene names, and gene count for each enriched term. Terms with adjusted P value ≤ 0.05 were considered significantly enriched. To infer potential biological functions associated with differentially expressed lncRNAs, protein-coding genes located within 100 kb upstream or downstream of each lncRNA were identified as putative cis-associated genes and used for GO and KEGG pathway enrichment analyses. GO enrichment was used to identify enriched biological processes, molecular functions, and cellular components, while KEGG enrichment was used to identify enriched pathways among the co-localized genes. Enrichment significance was assessed using adjusted P values, with adjusted P ≤ 0.05 considered significant.

Differential alternative splicing analysis was conducted using rMATS, a statistical framework for detecting alternative splicing changes from replicate RNA-seq datasets.^41^ rMATS identified alternative splicing events corresponding to skipped exons, mutually exclusive exons, alternative 5′ splice sites, alternative 3′ splice sites, and retained introns. Inclusion junction counts and skipping junction counts were calculated for each sample group, and biological replicates were represented as comma-separated values. Inclusion levels were calculated from normalized counts, and the inclusion level difference was defined as the average inclusion level of sample group 1 minus the average inclusion level of sample group 2. Differential splicing events were considered significant at an False Discovery Rate (FDR)-adjusted P value < 0.05.

### Fluorescent In Situ Hybridization (FISH) procedure

FISH was performed on mouse retinal sections using the RNAscope Assay protocol (Advanced Cell Diagnostics, Newark, CA, USA) according to the manufacturer’s instructions. Mice were perfused with PBS and 4% paraformaldehyde (PFA), eyes were extracted, enucleated, and post-fixed in PFA at 4 °C for 24 hours. Eyes were then cryoprotected in 30% sucrose overnight. Eyes were embedded in Tissue-Tek O.C.T. Compound (Sakura Finetek, Torrance, CA, USA) and cryosectioned at a thickness of 10 µm via cryostat. Retinal sections were mounted onto charged glass slides and stored at −80 °C until further processing. Prior to hybridization, slides were brought to room temperature, washed in PBS, baked at 60°C, fixed further in pre-chilled PFA, further dried with ethanol gradient treatment, a hydrophobic barrier was drawn around the sections using the ImmEdge Hydrophobic Barrier Pap Pen (Vector Laboratories, Newark, CA, USA) and subjected to target retrieval and protease treatment as recommended for frozen tissue sections. Sections were then incubated with RNAscope probes targeting *Prss56* (Mm-Prss56, Cat. No. 402701) and *Rbpms* (Mm-RBPMS-C2, Cat. No. 527231-C2) for SOHU samples, and *Lif* (Mm-Lif-C2, Cat. No. 475841), and *Rlbp1* (Mm-Rlbp1-C2, Cat. No. 468161-C2) for ONC samples. Hybridization was carried out in a humidified chamber at 40 °C using the proprietary hybridization system provided by ACD. Following probe hybridization, signal amplification was performed through a series of sequential amplification steps to generate detectable fluorescent signals. Fluorescent detection was achieved using tyramide signal amplification with Opal Fluorophores (Akoya Biosciences, Marlborough, MA, USA). Each fluorophore was applied at a dilution of 1:750. After completion of amplification and fluorophore incubations, sections were mounted and counterstained using VECTASHIELD Vibrance Antifade Mounting Medium (Vector Laboratories, Newark, CA, USA). Fluorescent imaging of retinal sections was performed using a BZ-X1000 fluorescence microscope (Keyence, Osaka, Japan) and a Nikon CSU-W1 SoRA confocal microscope (Nikon, Tokyo, Japan).

Quantification of *Lif* transcript signals was performed using ImageJ, where images were blinded before quantification. For each biological replicate, three retinal sections were analyzed as technical replicates, with n = 3 biological replicates per group. *Lif*-positive puncta were manually counted using the Cell Counter plugin within the inner nuclear layer (INL). Counts were obtained across the full INL length of each section and normalized to the measured INL length to calculate *Lif* puncta per millimeter. Values from the three technical replicate sections were averaged to generate one value per biological replicate for statistical analysis. Values were presented as mean ± SD. Because *Prss56* showed a robust and widespread increase in signal across the INL of SOHU retinas (Figure 8A and 8B), consistent with its significant upregulation identified by the RNA sequencing, FISH was used to confirm the direction and spatial distribution of expression rather than to estimate an additional quantitative effect size.

### Statistical tests

For Figure 8E, differences between ONC and SHAM groups were evaluated using an unpaired Welch’s t-test, with P < 0.05 considered statistically significant.

## RESULTS

Total RNA-seq generated an average of 63.5 million paired-end reads per ONC retina sample, with approximately 96% of reads mapping to the mouse genome. SOHU retina samples generated an average of approximately 61 million paired-end reads, with approximately 97% mapping to the mouse genome (*n* = 3 biological replicates per group). Initial PCA and unsupervised hierarchical clustering identified one discordant sample in each injury and corresponding sham condition for both the SOHU and ONC datasets. These samples were excluded from subsequent analyses, resulting in *n* = 2 biological replicates per group. PCA and heatmap clustering of the retained samples demonstrated clear separation between injury and corresponding sham groups (Figures 2A–C and 5A–C).

**Figure 2.**
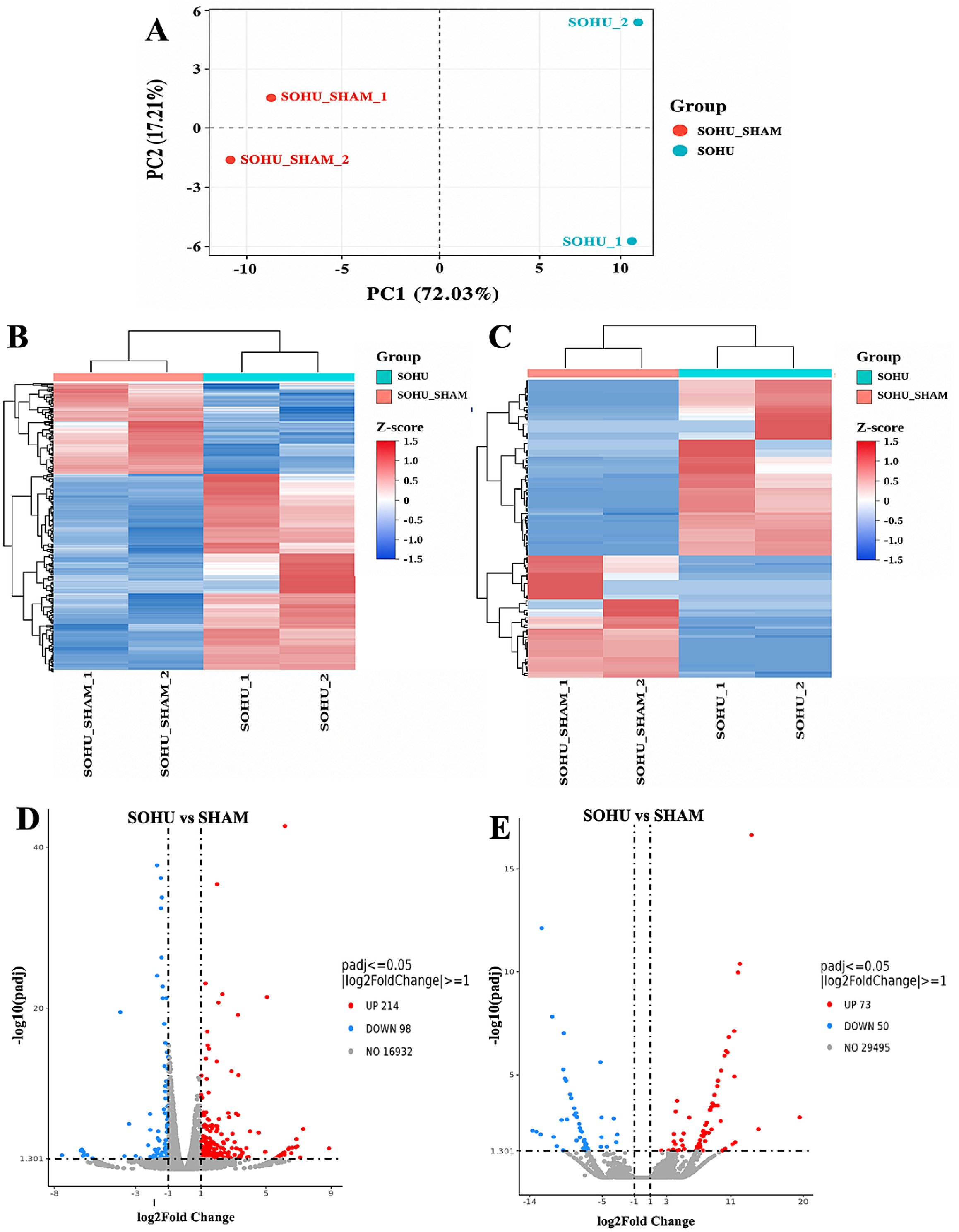
SOHU induction produces distinct retinal mRNA and lncRNA expression profiles. (**A**) Principal component analysis (PCA) of SOHU and SOHU_SHAM retinal RNA-seq samples showing separation between injury and sham groups. (**B**) Heat map showing hierarchical clustering of all 312 differentially expressed protein-coding genes identified in SOHU retinas compared with SOHU_SHAM controls. (**C**) Heat map showing hierarchical clustering of all 123 differentially expressed lncRNA-classified transcripts identified in SOHU retinas compared with SOHU_SHAM controls. Rows represent genes or transcripts and columns represent individual biological replicates. Red indicates relatively higher expression, and blue indicates relatively lower expression. (**D**) Volcano plot showing differentially expressed protein-coding genes in SOHU retinas compared with SOHU_SHAM controls. Red points indicate significantly upregulated genes, blue points indicate significantly downregulated genes, and gray points indicate transcripts that did not meet significance thresholds. (**E**) Volcano plot showing differentially expressed lncRNA-classified transcripts in SOHU retinas compared with SOHU_SHAM controls. n = 2 biological replicates per group for RNA-seq analysis. Differential expression was defined as absolute log2 fold change ≥ 1 and adjusted P value ≤ 0.05.

### rRNA-depleted total RNA-seq reveals coding and non-coding genes differentially expressed after SOHU induction

At 5 days post-SOHU induction, 312 coding genes were differentially expressed, including 214 upregulated and 98 downregulated coding genes (Figure 2D and Supplementary Table 1). A total of 123 lncRNAs were also differentially expressed, including 73 upregulated and 50 downregulated lncRNAs (Figure 2E and Supplementary Table 2). Differential expression analysis was performed using edgeR, with differentially expressed transcripts defined by an absolute log2 fold change ≥ 1 and adjusted p-value ≤ 0.05. Several top upregulated coding genes in SOHU retinas were associated with ocular hypertension, retinal stress, inflammatory signaling, or glaucoma-related biology, including *Prss56, Stat3, Edn2, C4b, Fgf2,* and *Serpina3n* ^42–47^. In addition, many differentially expressed lncRNAs were poorly characterized but showed robust changes in expression 5 days post-induction especially non-protein coding isoforms of protein coding genes (Figure 2E and Supplementary Table 2).

Functional enrichment analysis of significantly upregulated SOHU genes revealed a strong inflammatory and tissue-remodeling signature. GO terms were enriched for immune-response activation, leukocyte migration, extracellular matrix organization, basement membrane/collagen-associated categories, integrin binding, pattern-recognition receptor activity, and cell adhesion molecule binding (Figure 3A). KEGG analysis further supported enrichment of complement and coagulation cascades, phagosome, chemokine signaling, neutrophil extracellular trap formation, cell adhesion molecules, PI3K-Akt signaling, ECM-receptor interaction, and Fc gamma R-mediated phagocytosis (Figure 3B). Together, these results indicate that SOHU induction triggers early immune activation, extracellular matrix remodeling, cell-adhesion changes, and phagocytic/inflammatory signaling in whole retina.

**Figure 3.**
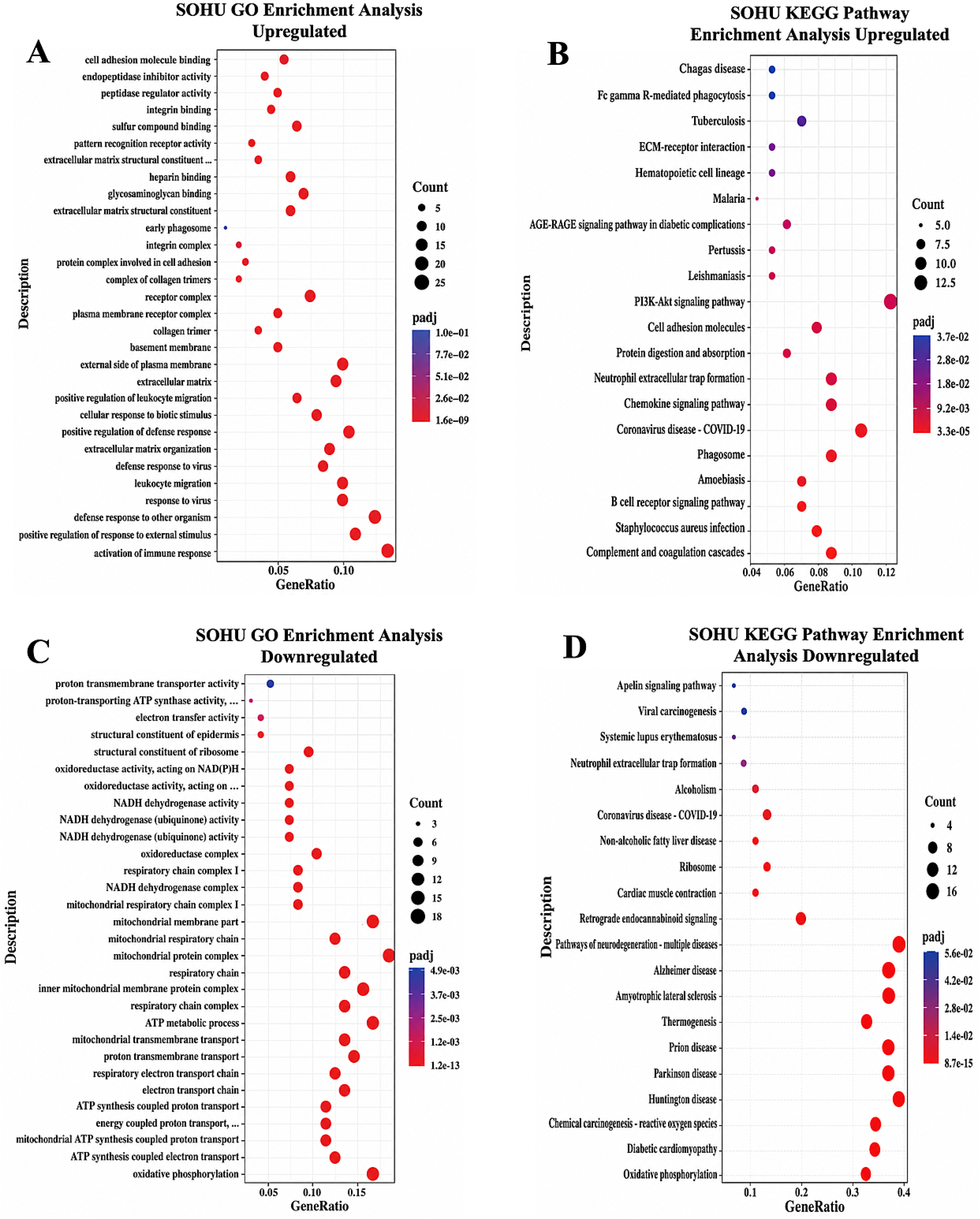
SOHU injury is associated with activation of inflammatory and extracellular matrix pathways and suppression of mitochondrial metabolic pathways. Functional enrichment analysis was performed using differentially expressed genes from SOHU retinas collected 5 days after ocular hypertension induction. **(A)** GO enrichment of upregulated genes showed prominent enrichment of immune activation, defense response, leukocyte migration, extracellular matrix organization, receptor complex, and cell adhesion-associated terms. **(B)** KEGG pathway analysis of upregulated genes identified enrichment of inflammatory and injury-associated pathways, including complement and coagulation cascades, phagosome, chemokine signaling pathway, cell adhesion molecules, PI3K–Akt signaling pathway, and ECM–receptor interaction. **(C)** GO enrichment of downregulated genes revealed strong enrichment of mitochondrial respiration and energy metabolism terms, including oxidative phosphorylation, ATP metabolic process, electron transport chain, respiratory chain, and mitochondrial respiratory chain complex. **(D)** KEGG pathway analysis of downregulated genes identified enrichment of oxidative phosphorylation, thermogenesis, and pathways associated with neurodegenerative disease. Dot size indicates gene count within each enriched term, while dot color indicates adjusted *P* value.

In contrast, enrichment analysis of significantly downregulated genes revealed suppression of mitochondrial and metabolic pathways. GO terms were dominated by oxidative phosphorylation, ATP synthesis-coupled electron transport, mitochondrial proton transport, electron transport chain, and respiratory chain complex/Complex I categories (Figure 3C). KEGG analysis similarly identified oxidative phosphorylation, reactive oxygen species-associated pathways, and neurodegeneration-related disease pathways (Figure 3D). Consistent with these enrichment results, the downregulated gene list was dominated by mitochondrial respiratory genes, including multiple mitochondrial NADH dehydrogenase subunits, mitochondrial cytochrome c oxidase subunits, mitochondrial ATP synthase subunits, and additional oxidative phosphorylation-associated genes such as *Ndufb8, Atp5e,* and *Cox6b2* (Supplementary Table 1). These findings suggest that SOHU induction is associated with reduced mitochondrial respiratory and energy metabolism gene expression, complementing the immune and extracellular matrix remodeling observed among upregulated genes.

Functional enrichment analysis of genes putatively cis-associated with differentially expressed lncRNAs identified pathways associated with immune regulation, autophagy, mitochondrial respiration, chromatin organization, metal and iron-sulfur cluster binding, necroptosis, AMPK/mTOR signaling, cell adhesion, neurotrophic signaling, and actin cytoskeleton regulation (Figure 4). These findings suggest that SOHU-responsive lncRNAs may participate in inflammatory, metabolic, and injury-related processes; however, the predicted regulatory relationships and biological functions of these lncRNAs require further experimental validation. Alternative splicing analysis was performed using rMATS to determine whether SOHU induction was associated with changes in transcript isoform expression in addition to overall gene expression. At 5 days post-SOHU induction, 320 significant differential alternative splicing events were identified, including 198 skipped exon events, 70 retained intron events, 23 alternative 5′ splice-site events, 18 alternative 3′ splice-site events, and 11 mutually exclusive exon events (Supplementary Figure 6 and Supplementary Table 5).

**Figure 4.**
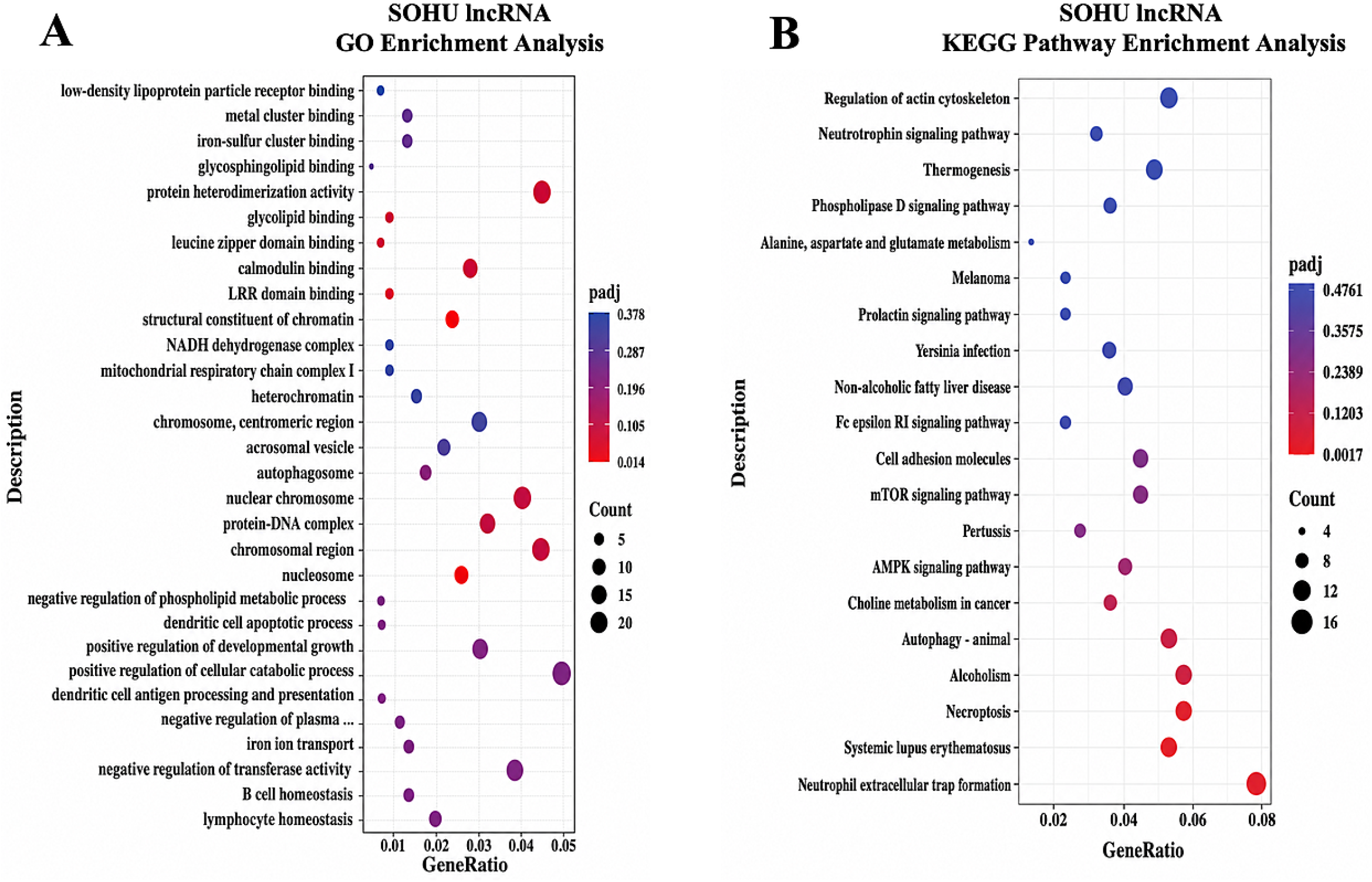
Differentially expressed lncRNAs in SOHU retinas are associated with genes involved in chromatin regulation, immune signaling, cell death, and stress-response pathways. Genomically proximal genes associated with differentially expressed lncRNAs in SOHU retinas 5 days after ocular hypertension induction were analyzed for GO and KEGG pathway enrichment. **(A)** GO enrichment analysis showed enrichment of terms related to chromatin and nuclear organization, protein heterodimerization, calmodulin binding, mitochondrial respiratory complex components, dendritic cell apoptotic processes, and immune cell homeostasis. **(B)** KEGG pathway analysis identified enrichment of injury- and stress-associated pathways, including neutrophil extracellular trap formation, necroptosis, autophagy, AMPK signaling, mTOR signaling, cell adhesion molecules, neurotrophic signaling, and regulation of the actin cytoskeleton. Dot size indicates the number of co-localized genes represented in each term, while dot color indicates the adjusted *P* value.

Protein-coding loci can give rise to multiple transcript isoforms through mechanisms such as alternative splicing, alternative promoter usage, and alternative transcriptional termination. While some of these isoforms encode proteins, others lack coding potential and are therefore classified as lncRNAs despite originating from the same genomic locus. Notably, a subset of SOHU-regulated lncRNA-classified transcripts corresponded to noncoding isoforms derived from annotated protein-coding genes (29 upregulated and 13 downregulated) (Supplementary Table 2). The potential implications of these isoform-level changes are further discussed in the Discussion section. Overall, these findings suggest that SOHU induction not only alters gene expression but also reshapes RNA processing and isoform usage, reflecting increased regulatory complexity in the early retinal injury response.

### Total RNA-seq reveals coding and non-coding genes differentially expressed 6 hours after ONC

PCA analyses and heatmap clustering demonstrate clear separation between sham and ONC injury groups (Figure 5A-5C). At 6-hours post-ONC, 301 coding genes were differentially expressed, including 217 upregulated and 84 downregulated coding genes (Figure 5D). Comparative analysis also identified 143 differentially expressed lncRNAs, including 110 upregulated and 33 downregulated lncRNAs (Figure 5E). Upregulated genes were dominated by early inflammatory, chemokine/cytokine, and vascular adhesion-associated responses, including *Ccl2, Ccl7, Cxcl10, Il1a, Il1rn, Icam1,* and *Sele* (Supplementary Table 3). Many ONC-regulated lncRNAs were poorly characterized, and a subset corresponded to non-protein-coding isoforms associated with annotated protein-coding genes (Supplementary Table 4).

**Figure 5.**
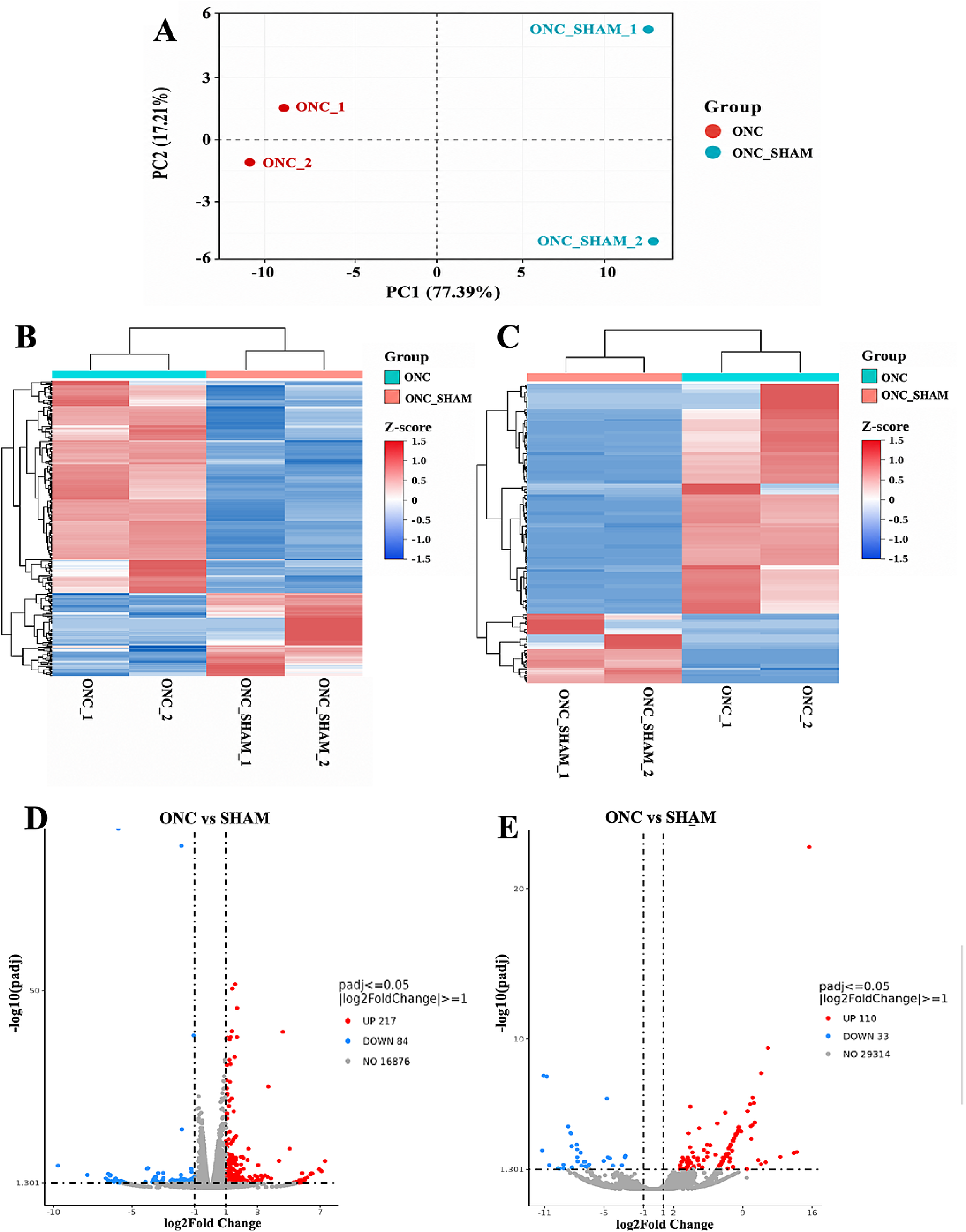
ONC exhibit distinct retinal mRNA and lncRNA expression profiles 6 hours after injury. **(A**) Principal component analysis (PCA) of ONC and ONC_SHAM retinal RNA-seq samples showing separation between injured and sham control groups. (**B**) Heat map showing hierarchical clustering of all 301 differentially expressed protein-coding genes identified in ONC retinas compared with ONC_SHAM controls. (**C**) Heat map showing hierarchical clustering of all 143 differentially expressed lncRNA-classified transcripts identified in ONC retinas compared with ONC_SHAM controls. Rows represent genes or transcripts and columns represent individual biological replicates. Red indicates relatively higher expression and blue indicates relatively lower expression. (**D**) Volcano plot showing differentially expressed protein-coding genes in ONC retinas compared with ONC_SHAM controls. Red points indicate significantly upregulated genes, blue points indicate significantly downregulated genes, and gray points indicate transcripts that did not meet significance thresholds. (**E**) Volcano plot showing differentially expressed lncRNA-classified transcripts in ONC retinas compared with ONC_SHAM controls. n = 2 biological replicates per group for RNA-seq analysis. Differential expression was defined as absolute log2 fold change ≥ 1 and adjusted P value ≤ 0.05.

Functional enrichment analysis of significantly upregulated genes revealed a rapid inflammatory and immune-associated retinal response. GO terms were enriched for chemokine activity, cytokine activity, chemokine receptor binding, leukocyte migration, neutrophil migration, granulocyte migration, and chemokine-mediated signaling (Figure 6A). KEGG analysis similarly identified enrichment of cytokine-cytokine receptor interaction, chemokine signaling, TNF signaling, IL-17 signaling, complement and coagulation cascades, NOD-like receptor signaling, and cell adhesion molecules (Figure 6B). Thus, these findings suggest that even at an acute 6-hour post-injury time point, ONC triggers a retinal inflammatory response characterized by chemokine/cytokine signaling, innate immune activation, and complement pathway involvement ^48–50^. In contrast, downregulated genes showed enrichment for ribosomal, intermediate filament/cytoskeletal, cell-junction, and keratin-associated terms (Figure 6C and 6D), although the strongest biologically coherent early-injury signal was observed among upregulated genes. Gene Ontology enrichment analysis of putative cis-associated protein-coding genes revealed terms related to glutathione metabolism, cytoskeletal organization, chromatin-associated structures, methylation, epigenetic regulation of gene expression, and neurotransmitter receptor regulatory activity (Figure 7A). KEGG enrichment analysis further identified pathways associated with glutathione metabolism, xenobiotic/drug metabolism, reactive oxygen species-related pathways, retrograde endocannabinoid signaling, Parkinson disease, amyotrophic lateral sclerosis, and retinol metabolism (Figure 7B). These results suggest that ONC-regulated lncRNAs may be associated with oxidative stress, metabolic regulation, chromatin/epigenetic regulation, cytoskeletal organization, and neurodegeneration-associated pathways; however, these associations are predictive and require experimental validation.

**Figure 6.**
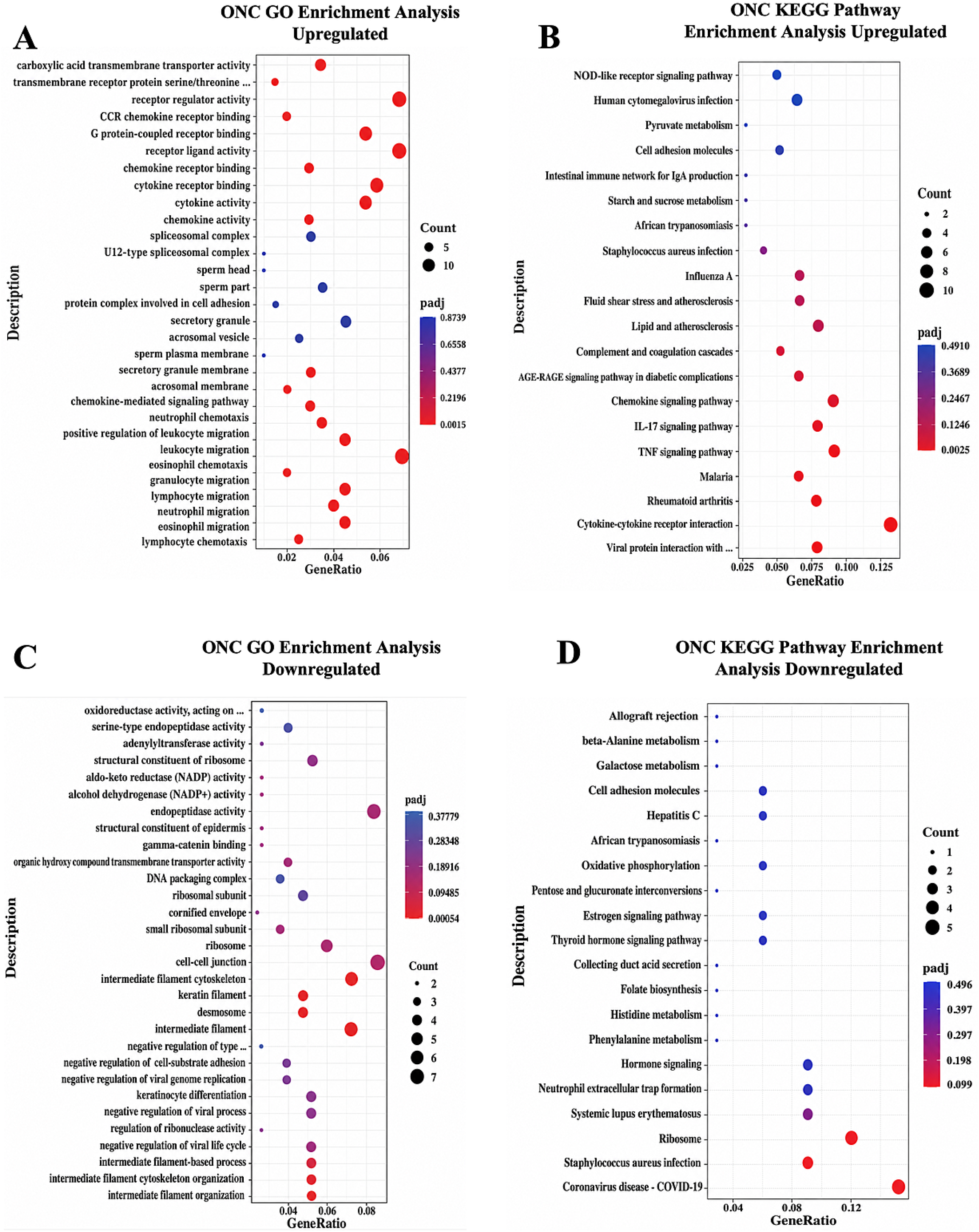
GO and KEGG enrichment analyses of ONC-regulated genes. Functional enrichment analysis was performed using differentially expressed genes from retinas collected 6 hours after optic nerve crush. **(A)** GO enrichment of upregulated genes showed prominent enrichment of chemokine activity, cytokine activity, chemokine receptor binding, chemokine-mediated signaling, leukocyte migration, neutrophil migration, granulocyte migration, and positive regulation of leukocyte migration. **(B)** KEGG pathway analysis of upregulated genes identified enrichment of inflammatory and immune-associated pathways, including cytokine-cytokine receptor interaction, chemokine signaling pathway, TNF signaling pathway, IL-17 signaling pathway, complement and coagulation cascades, NOD-like receptor signaling pathway, and cell adhesion molecules. **(C)** GO enrichment of downregulated genes revealed enrichment of ribosomal, intermediate filament/cytoskeletal, cell-junction, keratin-associated, and epidermal structural terms. **(D)** KEGG pathway analysis of downregulated genes identified enrichment of ribosome, oxidative phosphorylation, cell adhesion molecules, and additional immune/metabolic-associated pathways. Dot size indicates gene count within each enriched term, while dot color indicates adjusted P value.

**Figure 7.**
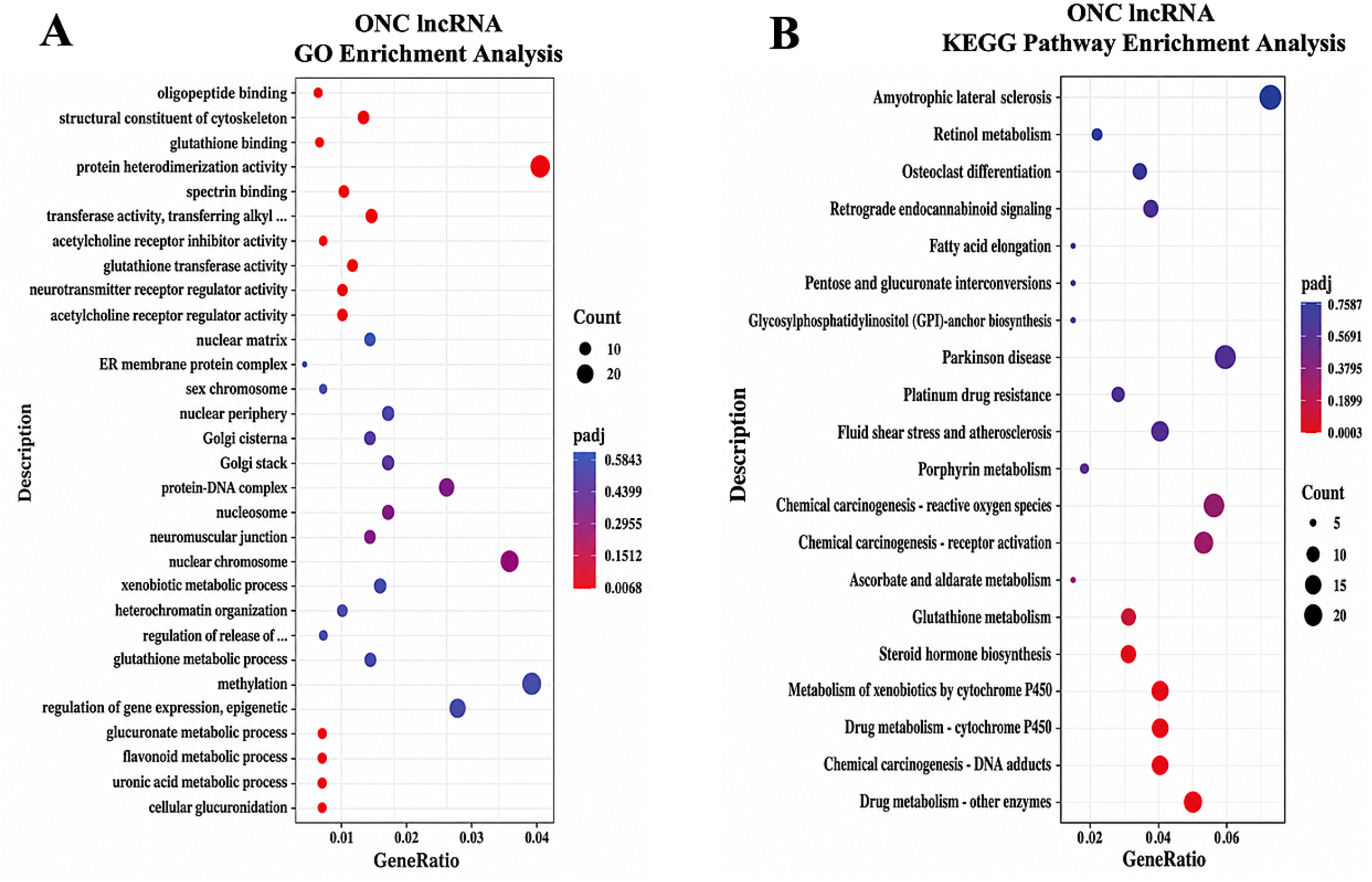
Functional enrichment analysis of genes putatively cis-associated with differentially expressed lncRNAs following ONC. Genomically proximal genes associated with differentially expressed lncRNAs in retinas collected 6 hours after optic nerve crush were analyzed for GO and KEGG pathway enrichment. **(A)** GO enrichment analysis showed enrichment of terms related to glutathione binding and glutathione metabolic processes, cytoskeletal organization, protein heterodimerization, chromatin and nuclear organization, methylation, epigenetic regulation of gene expression, xenobiotic metabolism, and neurotransmitter receptor regulatory activity. **(B)** KEGG pathway analysis identified enrichment of metabolic, oxidative stress, and neurodegeneration-associated pathways, including glutathione metabolism, drug/xenobiotic metabolism by cytochrome P450, chemical carcinogenesis-reactive oxygen species, retinol metabolism, retrograde endocannabinoid signaling, Parkinson disease, and amyotrophic lateral sclerosis. Dot size indicates the number of co-localized genes represented in each term, while dot color indicates the adjusted P value.

Because many ONC-regulated lncRNA-classified transcripts corresponded to non-protein-coding transcript isoforms arising from annotated protein-coding genes, we further examined alternative splicing patterns in the ONC RNA-seq dataset to assess transcript-level changes associated with ONC. At 6-hours post-ONC, 452 significant differential alternative splicing events were identified, including 198 retained intron events, 178 skipped exon events, 37 alternative 5′ splice-site events, 22 alternative 3′ splice-site events, and 17 mutually exclusive exon events (Supplementary Figure 6 and Supplementary Table 6). These findings provide additional evidence of widespread changes in transcript structure and splicing patterns following ONC, complementing the lncRNA analysis, which identified 69 ONC-regulated lncRNA-classified transcripts associated with annotated protein-coding genes, including 60 upregulated and 9 downregulated transcripts (Supplementary Table 4). Together, these results suggest that ONC induces changes at multiple levels of transcript regulation, including differential expression of non-protein-coding transcript isoforms and alternative splicing, although the specific relationship between individual lncRNA-classified transcripts and differential splicing events remains to be determined.

### FISH Validation of Select DEGs

We selected serine protease 56 (*Prss56*) for expression validation based on its known association with angle-closure glaucoma-related ocular phenotypes ^44^ and its strong upregulation following SOHU induction (Figure 8A and 8B). FISH qualitatively confirmed a marked increase in *Prss56* transcript signal in SOHU retinas compared with the SHAM controls. The increased signal was evident in representative retinal sections and was broadly distributed across the retina (Figure 8B), consistent with the direction of differential expression identified by RNA sequencing. Additionally, *Lif* was selected for validation based on its robust upregulation in the ONC RNA-seq dataset and prior evidence implicating LIF in Müller glial activation following optic nerve injury and in retinal stress-responsive signaling.^51,52^ FISH demonstrated increased *Lif* transcript signal in ONC retinas compared with SHAM controls, with signal detected primarily within the INL (Figure 8C and 8D). Notably, the increase in *Lif* expression was predominantly observed in the retinal regions adjacent to the optic nerve head, but not in the peripheral retina (Figure 8D). This spatially restricted *Lif* expression suggests that injury-induced signaling following ONC may propagate from the injury site into the proximal retina but does not uniformly spread throughout the entire retinal tissue at this early time point. Quantification of *Lif*-positive puncta confirmed a significant increase in ONC retinas relative to SHAM controls (Figure 8E). Together, these findings support the RNA-seq results and independently validate the injury-associated upregulation of *Prss56* in the SOHU model and *Lif* following ONC.

**Figure 8.**
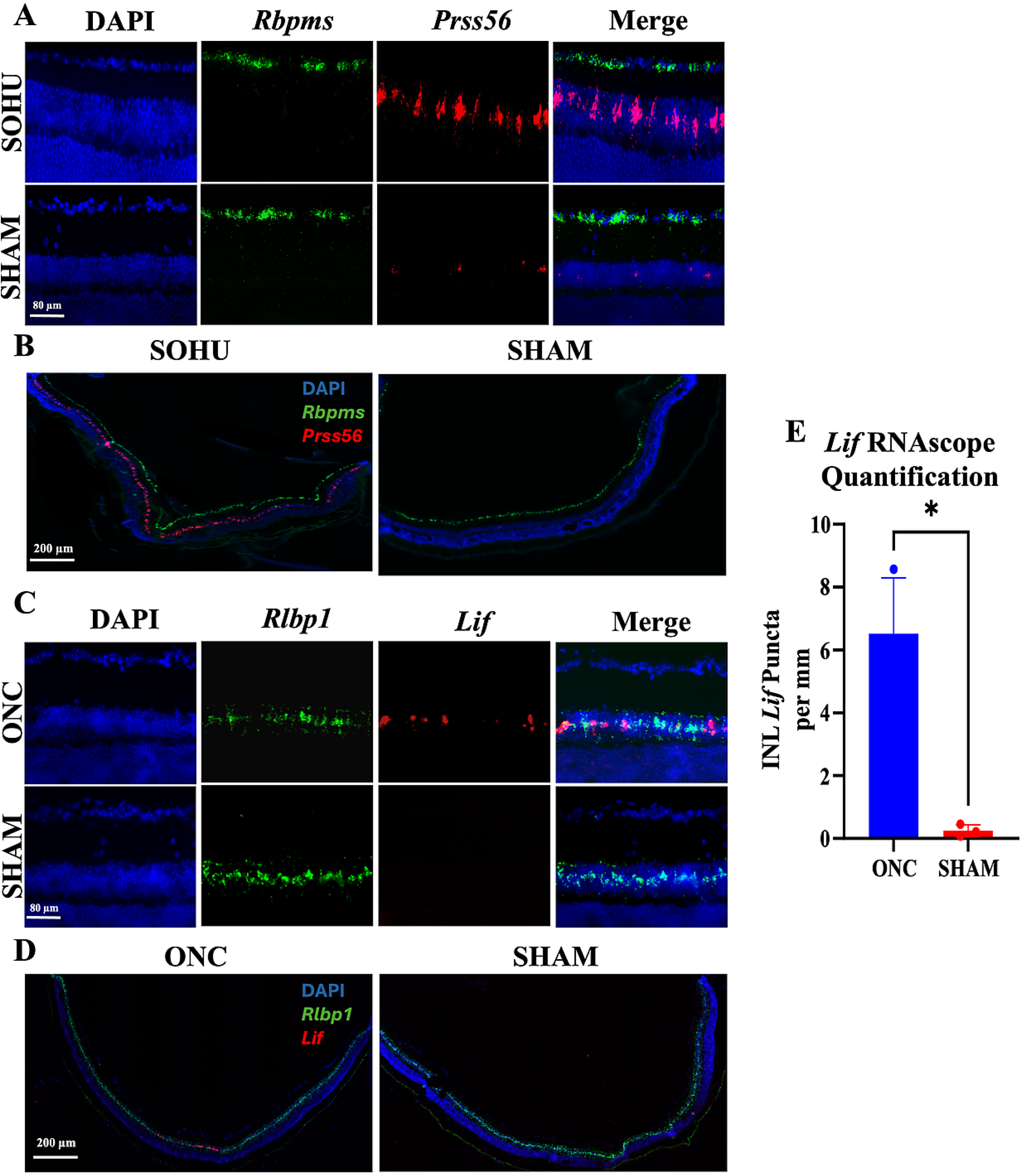
FISH validation of *Prss56* and *Lif* expression in experimental retinal injury models. **(A)** 40x and **(B)** 20x magnification representative images for *Prss56* expression in retinal sections from SOHU and SHAM eyes. Sections were counterstained with DAPI and co-labeled with *Rbpms*. *Prss56* transcript signal was markedly increased in SOHU retinas compared with SHAM controls, supporting the RNA-seq identification of *Prss56* as a strongly upregulated SOHU-associated transcript. **(C)** 40x and **(D)** 20x magnification representative images for *Lif* expression in retinal sections from ONC and SHAM eyes. Sections were counterstained with DAPI and co-labeled with retinaldehyde-binding protein 1 (*Rlbp1*) to identify Müller cells. *Lif* transcript signal is increased in ONC retinas compared with SHAM controls and is detected within the inner nuclear layer (INL) in association with *Rlbp1*-positive regions. **(E)** Quantification of *Lif*-positive puncta within the INL showed significantly greater *Lif* transcript signal in ONC retinas than in SHAM controls. n = 3 retinas per group. Data are presented as mean ± SD. *P* < 0.05 by unpaired Welch’s *t*-test. Scale bars = 80 µm for 40x and 200 µm for 20x

## DISCUSSION

In this study, we sought to identify early coding and noncoding transcriptional changes in two optic nerve injury models, SOHU and ONC. Although SOHU and ONC likely differ in both injury mechanisms and different tissue collection time points were used respectively, each exhibited significant transcriptomic alterations in the whole retina. Both models showed differential expression of protein-coding genes and lncRNA-classified transcripts, along with injury-associated alternative splicing events. SOHU retinas demonstrated prominent changes in immune signaling, extracellular matrix remodeling, complement activation, phagocytic pathways, and mitochondrial/metabolic processes, whereas ONC retinas mounted a rapid chemokine/cytokine-mediated innate immune response as early as 6 hours after injury. Collectively, these findings indicate that early retinal injury is characterized not only by the activation of canonical injury-response genes but also by widespread alterations in noncoding RNA expression and RNA processing.

In the SOHU model, the most prominent protein-coding gene changes suggested activation of inflammatory, stress-response, complement, and tissue-remodeling pathways. Among the most strongly upregulated genes were *Prss56, Edn2, Fgf2, C4b, Serpina3n, Stat3, and Serpine3*, which have reported relevance to ocular hypertension, retinal injury, or glaucoma-associated pathology ^42–47,53^. Validation of *Prss56* upregulation supports the reliability of the RNA-seq dataset and suggests that SOHU induction produces reproducible transcriptional changes relevant to ocular hypertension-associated retinal stress. In addition, upregulation of *C4b* is consistent with early complement-cascade activation in experimental glaucoma, where classical complement signaling has been implicated in RGC dendritic and synaptic degeneration^54,55^, while increased *Fgf2* and *Stat3* expression may reflect activation of glial, cytokine-responsive, or injury-associated survival pathways.^56–59^ *Serpine3* and *Serpina3n* upregulation further suggest changes in protease inhibitor activity, extracellular matrix regulation, and injury-associated tissue remodeling.^47,53^ Consistent with these gene-level findings, GO and KEGG enrichment analyses identified immune activation, leukocyte migration, complement/coagulation signaling, extracellular matrix organization, basement membrane/collagen-associated terms, integrin binding, and cell adhesion pathways. Together, these findings support a model in which SOHU induction produces an early whole-retina response involving inflammatory signaling, complement activation, retinal stress-associated gene expression, and extracellular matrix remodeling.

In contrast to the inflammatory and extracellular matrix-associated pathways enriched among upregulated genes, downregulated SOHU genes were strongly associated with mitochondrial oxidative phosphorylation and respiratory chain function, including electron transport chain activity, NADH dehydrogenase activity, proton transport, and ATP synthesis-associated terms. This pattern suggests that early ocular hypertension-associated retinal injury may involve reduced expression of mitochondrial energy metabolism genes. Because the RNA-seq was performed on whole retina, these changes may reflect altered metabolic activity across multiple retinal cell types rather than a single cell population. The enrichment of neurodegeneration-related KEGG pathways should therefore be interpreted cautiously, as these categories often share mitochondrial and oxidative phosphorylation gene sets rather than indicating disease-specific mechanisms.

Among the differentially expressed lncRNAs identified in SOHU retinas, we observed that multiple *Gas5* transcript isoforms were significantly upregulated and a significant retained intron event was also detected at the *Gas5* locus. These findings suggest that SOHU injury may regulate *Gas5* at both the expression and transcript-processing levels. Given prior evidence implicating Gas5 in RGC loss in a rat glaucoma model,^60^ our results highlight *Gas5* as a strong candidate lncRNA for further investigation using the SOHU model. However, the functional consequence of *Gas5* upregulation and altered splicing in the SOHU model remains unknown and will require targeted validation and perturbation experiments.

Consistent with this broader pattern, many of the differentially expressed lncRNA-classified transcripts identified in SOHU retinas may represent alternative noncoding transcript variants associated with annotated protein-coding loci. This observation suggests that SOHU injury induces transcript-level remodeling in which individual gene loci may alter their isoform output in addition to changing overall gene expression. Such changes highlight an additional regulatory layer beyond canonical differential gene expression. Functionally, noncoding isoforms derived from protein-coding loci may influence gene expression, RNA stability, translation, or serve as regulatory decoys or scaffolds; however, these potential functions remain to be experimentally tested in the context of ocular hypertension-associated retinal injury. In addition to *Gas5*, *LINC54*-annotated transcripts emerged as shared injury-associated lncRNA candidates across both retinal injury models (Supplementary Tables 2 and 4). A *LINC54* transcript was strongly upregulated in SOHU retinas, and a *LINC54*-annotated transcript was also significantly upregulated 6 hours after ONC. Although these transcripts are currently poorly characterized and require further annotation, their induction in both ocular hypertension-associated and acute optic nerve injury contexts suggests that *LINC54*-associated noncoding transcription may represent a shared feature of the retinal injury response.

The ONC dataset provided an acute optic nerve injury comparison and revealed a rapid inflammatory transcriptional response by 6 hours post-ONC. Upregulated genes were dominated by chemokines, cytokines, and adhesion-associated molecules, including *Ccl2, Ccl7, Cxcl10, Il1a, Il1rn, Icam1, Sele*, and *Lif*. Validation of *Lif* upregulation supports the reliability of the ONC RNA-seq dataset and suggests that ONC induces reproducible transcriptional changes related to injury-associated cytokine and retinal stress-response signaling.

The robust induction of chemokine, cytokine, and inflammatory signaling pathways as early as 6 hours after axonal injury was somewhat unexpected, given that well-established RGC injury-responses including the activation of endothelin converting enzyme like 1(Ecel1) and induction of Atf3 and Atf4 are typically reported to increase more prominently at later time points, including approximately 1 day after ONC.^17,61^ The early appearance of inflammatory signaling therefore suggests that acute axonal injury triggers retinal responses that precede, or occur independently of, the canonical RGC stress-response program. In this regard, it is noteworthy that *Lif* expression, as detected by FISH, was predominantly localized to the INL. To identify Müller cells, we performed RNAscope using a probe against *Rlbp1*, a well-established Müller glial marker.^62^ Publicly available adult mouse retinal scRNA-seq data, together with prior experimental evidence, indicate that *Lif* expression is enriched in Müller glia in the intact retina.^52,63^ However, our FISH results revealed that following ONC, *Lif* expression was not restricted to Müller cells. Although some *Lif*-positive cells co-expressed *Rlbp1*, additional *Lif* expression was observed in other retinal cell types. The cellular sources and initiating mechanisms underlying this early cytokine induction remain unclear. However, the spatial distribution of *Lif* expression may provide clues. At 6 hours post-ONC, *Lif* mRNA was observed predominantly within the INL and was particularly enriched in retinal regions adjacent to the optic nerve head. This pattern raises the possibility that acute axonal injury generates a rapidly propagated injury signal from the site of optic nerve damage to nearby retinal cells, including Müller glia and other INL-resident cell types. Such signaling could be mediated by retrograde stress signals from injured axons, alterations in axon-glia communication, or local mechanical and metabolic disturbances at the optic nerve head.

Consistent with these gene-level changes, GO and KEGG enrichment analyses identified chemokine activity, cytokine signaling, leukocyte migration, TNF signaling, IL-17 signaling, complement/coagulation, and cell adhesion pathways. Differences in enriched pathways between SOHU and ONC likely reflect the distinct injury mechanisms and time points examined, with ONC capturing an acute axonal injury response at 6 hours and SOHU capturing an early ocular hypertension-associated retinal stress response at 5 days. These respective time points were selected to capture early retinal injury-associated transcriptional responses before substantial RGC soma loss would be expected based on reported degeneration time courses in ONC and SOHU models. ^28–30,64^ Therefore, the transcriptional changes observed in both models are best interpreted as early whole-retina stress, inflammatory, metabolic, and RNA-processing responses rather than changes driven primarily by RGC depletion. Nevertheless, both datasets revealed inflammatory pathway activation, lncRNA dysregulation, and alternative splicing changes, suggesting that protein-coding, noncoding, and isoform-level remodeling may represent shared features of early retinal injury.

The lncRNA-associated pathway analyses further suggest that injury-regulated lncRNAs may participate in retinal stress responses. In SOHU retinas, putative cis-associated genes were linked to immune regulation, autophagy, mitochondrial respiratory function, chromatin organization, cell adhesion, necroptosis, and neurotrophin-related signaling. This pattern is consistent with the broader SOHU transcriptional profile, which showed immune activation, extracellular matrix remodeling, complement/phagocytic signaling, and reduced mitochondrial energy metabolism. In contrast, ONC-associated lncRNA target pathways were more strongly linked to oxidative stress and detoxification-related metabolism, including glutathione, glucuronidation, xenobiotic metabolism, chromatin/epigenetic regulation, cytoskeletal organization, and neuronal signaling-related pathways. Although these differences may reflect distinct injury mechanisms and collection time points, both models support the idea that lncRNA expression changes are coupled to broader retinal stress and injury-response pathways. Importantly, these analyses remain hypothesis-generating because genomic proximity does not establish direct regulatory interactions between lncRNAs and nearby protein-coding genes.

In addition to differential gene and lncRNA expression, both injury models showed evidence of altered transcript isoform usage. These findings suggest that early retinal injury is associated with altered RNA processing and transcript-level remodeling. This is particularly relevant to the lncRNA analysis, as many injury-regulated lncRNA-classified transcripts corresponded to noncoding isoforms associated with annotated protein-coding loci. Although the functional significance of these isoforms remains unknown, this observation is consistent with the concept that cellular stress and injury can alter RNA-processing programs, including alternative splicing, intron retention, and production of nonproductive or noncoding transcript variants.^65–67^ Intron retention and alternative splicing have been shown to regulate mammalian gene-expression programs,^68,69^ and alternative splicing can generate transcript isoforms that are not translated into canonical proteins but may instead undergo nuclear retention, degradation through RNA-surveillance pathways such as nonsense-mediated decay, or other forms of post-transcriptional regulation.^70–72^ Therefore, the substantial proportion of noncoding isoform-like lncRNA candidates observed in SOHU and ONC may reflect injury-associated remodeling of transcript processing rather than regulation of only canonical intergenic lncRNAs. Future work using long-read sequencing, isoform-specific PCR, or targeted validation will be needed to determine whether these injury-regulated isoforms are stable, cell-type-specific, and functionally relevant in retinal injury.

Comparison of SOHU and ONC revealed both shared and model-specific features of retinal injury. ONC produced a rapid chemokine/cytokine-dominant response, whereas SOHU showed a broader ocular hypertension-associated profile characterized by immune activation, extracellular matrix remodeling, complement/phagocytic signaling, and reduced mitochondrial energy metabolism gene expression. The two models also differed at the transcript-processing level, with ONC exhibiting a greater number of significant alternative splicing events than SOHU. This increase may reflect the abrupt nature of acute axonal injury in ONC compared with the more gradual stress response induced by SOHU,^28,73^ although additional time-course studies would be required to determine whether these differences are driven by injury kinetics, injury mechanism, or both. Retained intron events were particularly prominent in ONC, which may indicate rapid stress-associated regulation of transcript maturation, nuclear retention, or RNA surveillance pathways. In contrast, skipped exon events were the most frequent category in SOHU, potentially reflecting injury-associated isoform switching during ocular hypertension-associated retinal stress.^74^ Despite these model-specific differences, both datasets showed differential lncRNA expression and altered isoforms, suggesting that noncoding RNA regulation and transcript remodeling may represent common features of early retinal injury responses.

### Limitations of the study

This study has several limitations. First, RNA-seq was performed with a limited number of biological replicates, and additional samples would strengthen the differential expression and alternative splicing analyses. Second, whole-retina RNA-seq does not identify the specific retinal cell types responsible for the observed transcriptional changes. Because RGCs, Müller glia, microglia, astrocytes, vascular cells, and other retinal cell populations may all contribute to the bulk RNA signal, future single-cell or spatial transcriptomic approaches will be needed to resolve cell-type-specific responses. Third, proximity-based identification of putative cis-associated genes is predictive and does not establish direct regulatory interactions between lncRNAs and nearby protein-coding genes. Fourth, because only male mice were included, potential sex-dependent differences in retinal injury-associated transcriptional responses were not evaluated. Finally, while FISH validation confirmed expression changes for two representative genes, *Prss56* and *Lif*, validation was not performed for the broader set of differentially expressed transcripts including lncRNAs. Future studies could validate additional candidates using FISH; however, multiplexed spatial transcriptomic methods such as Multiplexed Error-Robust Fluorescence In Situ Hybridization (MERFISH)^75^ may offer greater insight by simultaneously determining the magnitude and spatial distribution of gene expression changes across retinal cell types. Our dataset provides a resource for prioritizing candidate genes for these future spatial investigations.

Overall, this study identifies early coding, noncoding, and alternative splicing changes in SOHU and ONC retinas. The datasets generated here provide a resource for future mechanistic studies aimed at defining the cellular origins, functional significance, and regulatory roles of injury-associated coding and noncoding transcripts in the retina.

## Funding Information

This work was supported by grants from the National Eye Institute (NEI) R01EY032542 (K.K.P), R01EY034531 (K.K.P), P30EY030413 (K.K.P), Startup funds from the UT Southwestern Department of Ophthalmology, The Fichtenbaum Charitable Trust Research Award, Anne Marie & Thomas B. Walker Fund, and Challenge Grant from Research to Prevent Blindness (K.K.P).

## Commercial Relationships Disclosure

None

## Notes

### Competing Interest Statement

The authors have declared no competing interest.

