## Supplementary material for "Total RNA Sequencing Reveals Early Coding, Noncoding, and Isoform-Level Transcriptomic Changes Following Traumatic and Glaucomatous Optic Nerve Injury": Sup Figs.Legends

**Supplementary Figure 1. Classification of raw RNA-seq reads following quality filtering in ONC and SOHU samples.** Pie charts show the proportions and numbers of clean reads, reads containing undetermined bases (N), low-quality reads, and adapter-related reads for retained ONC, ONC_SHAM, SOHU, and SOHU_SHAM biological replicates. Most samples retained more than 99% clean reads after filtering. SOHU_SHAM_1 retained 91.14% clean reads, with 8.85% of reads classified as adapter-related. Reads containing adapters, reads containing more than 0.1% undetermined bases, and reads in which more than 50% of bases had a quality score of 5 or lower were removed before downstream analysis.

**Supplementary Figure 2. RNA-seq quality control metrics for ONC and SOHU samples.** RNA-seq quality-control plots are shown for retained ONC, ONC_SHAM, SOHU, and SOHU_SHAM biological replicates. (**A**) Sequencing error-rate distributions across read positions. (**B**) Nucleotide-content distributions across read positions, showing the percentages of adenine, thymine, guanine, cytosine, and undetermined bases.

**Supplementary Figure 3. Genomic region distribution of mapped RNA-seq reads.** Pie charts show the proportion of reads mapping to exonic, intronic, and intergenic regions for each biological replicate. Across samples, a majority of mapped reads aligned to exonic regions, with smaller proportions mapping to intronic and intergenic regions, supporting successful alignment of RNA-seq reads to annotated genomic features prior to downstream differential expression, lncRNA, enrichment, and alternative splicing analyses.

**Supplementary Figure 4. Characterization of identified lncRNA transcripts in ONC and SOHU RNA-seq datasets.** Candidate lncRNA transcripts were identified from assembled RNA-seq transcripts and evaluated using coding-potential and transcript feature analyses. **(A)** Venn diagrams showing candidate lncRNA filtering by Coding Potential Calculator (CPC), Coding-

Non-Coding Index (CNCI), and Protein family database (PFAM) in ONC and SOHU datasets. **(B)** Exon number density distributions comparing annotated lncRNAs, annotated mRNAs, novel lncRNAs, and novel mRNAs in ONC and SOHU datasets. **(C)** Transcript length density distributions comparing annotated and novel lncRNA/mRNA transcripts. **(D)** Open reading frame (ORF) length density distributions comparing annotated and novel lncRNA/mRNA transcripts. These analyses support lncRNA classification by showing expected differences in coding potential and transcript features between lncRNAs and mRNAs.

**Supplementary Figure 5. FPKM expression distribution across ONC and SOHU RNA-seq samples.**

Boxplots showing the distribution of log10(FPKM + 1) expression values across biological replicates in the ONC and SOHU RNA-seq datasets. FPKM values were used to estimate gene/transcript expression while accounting for sequencing depth and transcript length. Similar expression distributions across injury and sham samples support comparable global expression profiles prior to downstream differential expression, lncRNA, enrichment, and alternative splicing analyses.

**Supplementary Figure 6. Differential alternative splicing events identified in ONC and SOHU retinas.**

Alternative splicing analysis was performed using rMATS to identify significant differential splicing events in ONC and SOHU RNA-seq datasets. Bar graph shows the number of significant alternative splicing events detected in each category, including skipped exon (SE), retained intron (RI), alternative 5′ splice site (A5SS), alternative 3′ splice site (A3SS), and mutually exclusive exon (MXE) events. ONC retinas exhibited 452 significant differential splicing events, while SOHU retinas exhibited 320 significant differential splicing events.

Retained intron and skipped exon events represented the most frequent categories across the two injury models.

**Supplementary Table 1. Differentially expressed protein-coding genes in SOHU retinas.** This table includes all significantly upregulated and downregulated protein-coding genes identified in SOHU retinas compared with SOHU_SHAM controls. Differential expression was defined as absolute log2 fold change ≥ 1 and adjusted p-value ≤ 0.05. Separate sheets list upregulated and downregulated genes and include gene ID, gene name, gene description, genomic locus, normalized expression values for individual biological replicates, group mean expression values, log2 fold change, p-value, and adjusted p-value.

**Supplementary Table 2. Differentially Expressed Noncoding/lncRNA-Classified Transcripts in SOHU Retinas.** This table includes significantly upregulated and downregulated noncoding/lncRNA-classified transcripts identified in SOHU retinas compared with SOHU_SHAM controls. Differential expression was defined as absolute log2 fold change ≥ 1 and adjusted p-value ≤ 0.05. Transcripts were grouped into broad annotation-based categories using transcript annotation, gene/locus name, genomic context, and gene description information. Categories include noncoding isoforms of protein-coding loci, RIKEN/predicted/novel loci, intronic/overlapping/antisense transcripts, known lncRNA/host-gene loci, and LINC/novel intergenic lncRNAs. These categories are intended to summarize annotation patterns and do not indicate experimentally validated transcript function.

**Supplementary Table 3. Differentially Expressed Protein-Coding Genes in ONC Retinas.** This table includes all significantly upregulated and downregulated protein-coding genes identified in ONC retinas compared with ONC_SHAM controls. Differential expression was defined as absolute log2 fold change ≥ 1 and adjusted p-value ≤ 0.05. Separate sheets list upregulated and downregulated genes and include gene ID, gene name, gene description, genomic locus, normalized expression values for individual biological replicates, group mean expression values, log2 fold change, p-value, and adjusted p-value.

**Supplementary Table 4. Differentially Expressed Noncoding/lncRNA-Classified Transcripts in ONC Retinas.** This table includes significantly upregulated and downregulated noncoding/lncRNA-classified transcripts identified in ONC retinas compared with ONC_SHAM controls. Differential expression was defined as absolute log2 fold change ≥ 1 and adjusted p-value ≤ 0.05. Transcripts were grouped into broad annotation-based categories using transcript annotation, gene/locus name, genomic context, and gene description information. Categories include noncoding isoforms of protein-coding loci, RIKEN/predicted/novel loci, intronic/overlapping/antisense transcripts, known lncRNA/host-gene loci, and LINC/novel intergenic lncRNAs. These categories are intended to summarize annotation patterns and do not indicate experimentally validated transcript function.

**Supplementary Table 5. Significant Differential Alternative Splicing Events Identified in SOHU Retinas.** This table includes significant differential alternative splicing events identified in SOHU retinas compared with SOHU_SHAM controls using rMATS. Alternative splicing events are organized into separate sheets by event type, including alternative 3′ splice site (A3SS), alternative 5′ splice site (A5SS), mutually exclusive exon (MXE), retained intron (RI), and skipped exon (SE) events. For each event, the table includes gene ID, gene symbol, chromosome, strand, event-specific genomic coordinates, inclusion junction counts, skipping junction counts, inclusion and skipping form lengths, p-value, false discovery rate (FDR), replicate-level inclusion values, and inclusion level difference. Inclusion level difference was calculated as the average inclusion level of sample group 1 minus the average inclusion level of sample group 2. Differential alternative-splicing events were considered significant at FDR < 0.05.

**Supplementary Table 6. Significant Differential Alternative Splicing Events Identified in ONC Retinas.** This table includes significant differential alternative splicing events identified in ONC retinas compared with ONC_SHAM controls using rMATS. Alternative splicing events are organized into separate sheets by event type, including alternative 3′ splice site (A3SS), alternative 5′ splice site (A5SS), mutually exclusive exon (MXE), retained intron (RI), and skipped exon (SE) events. For each event, the table includes gene ID, gene symbol, chromosome, strand, event-specific genomic coordinates, inclusion junction counts, skipping junction counts, inclusion and skipping form lengths, p-value, false discovery rate (FDR), replicate-level inclusion values, and inclusion level difference. Inclusion level difference was calculated as the average inclusion level of sample group 1 minus the average inclusion level of sample group 2. Differential alternative-splicing events were considered significant at FDR < 0.05.
