## Supplementary material for "Total RNA Sequencing Reveals Early Coding, Noncoding, and Isoform-Level Transcriptomic Changes Following Traumatic and Glaucomatous Optic Nerve Injury": Sup.Figs

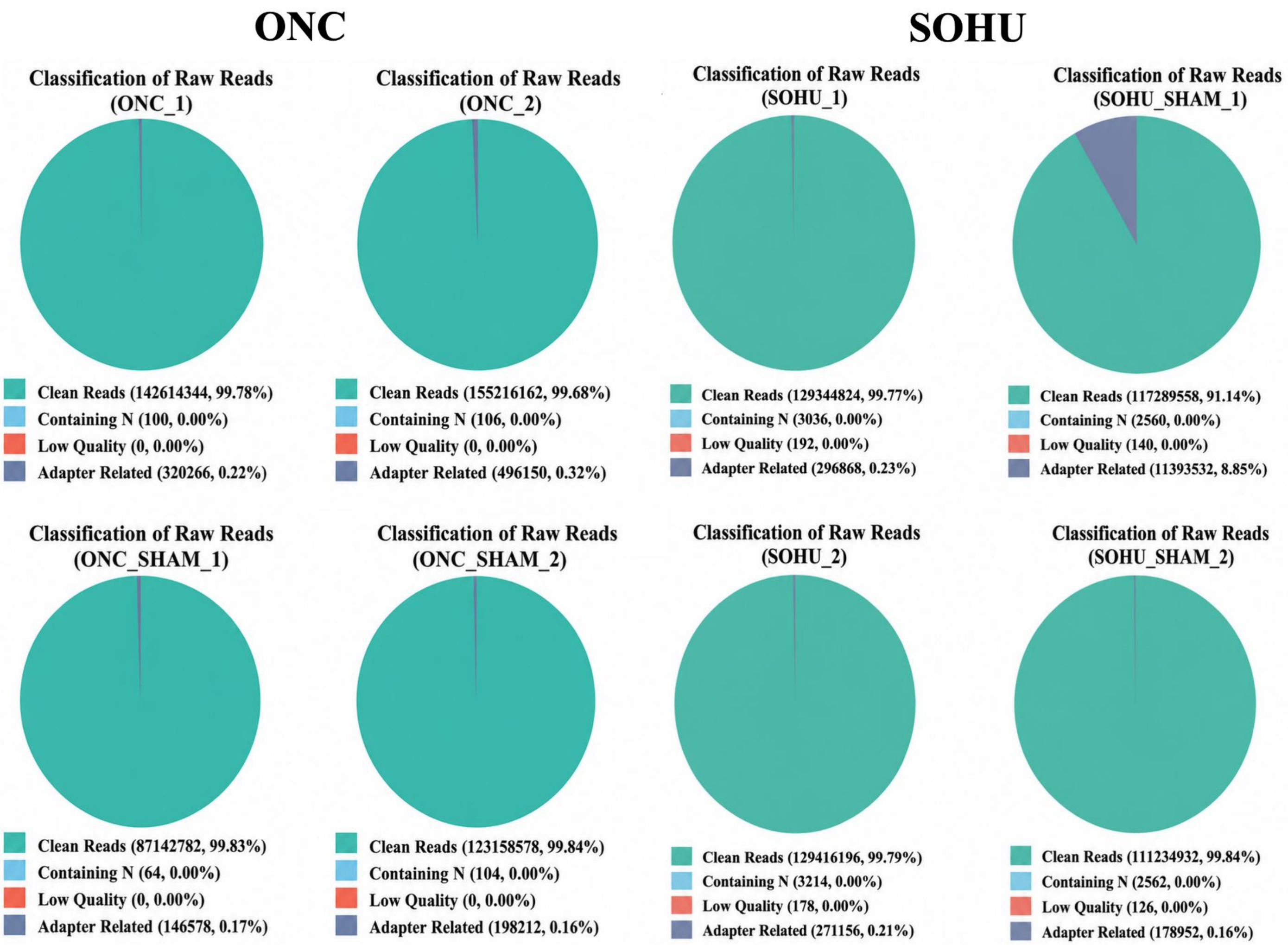


Supplementary Figure 1

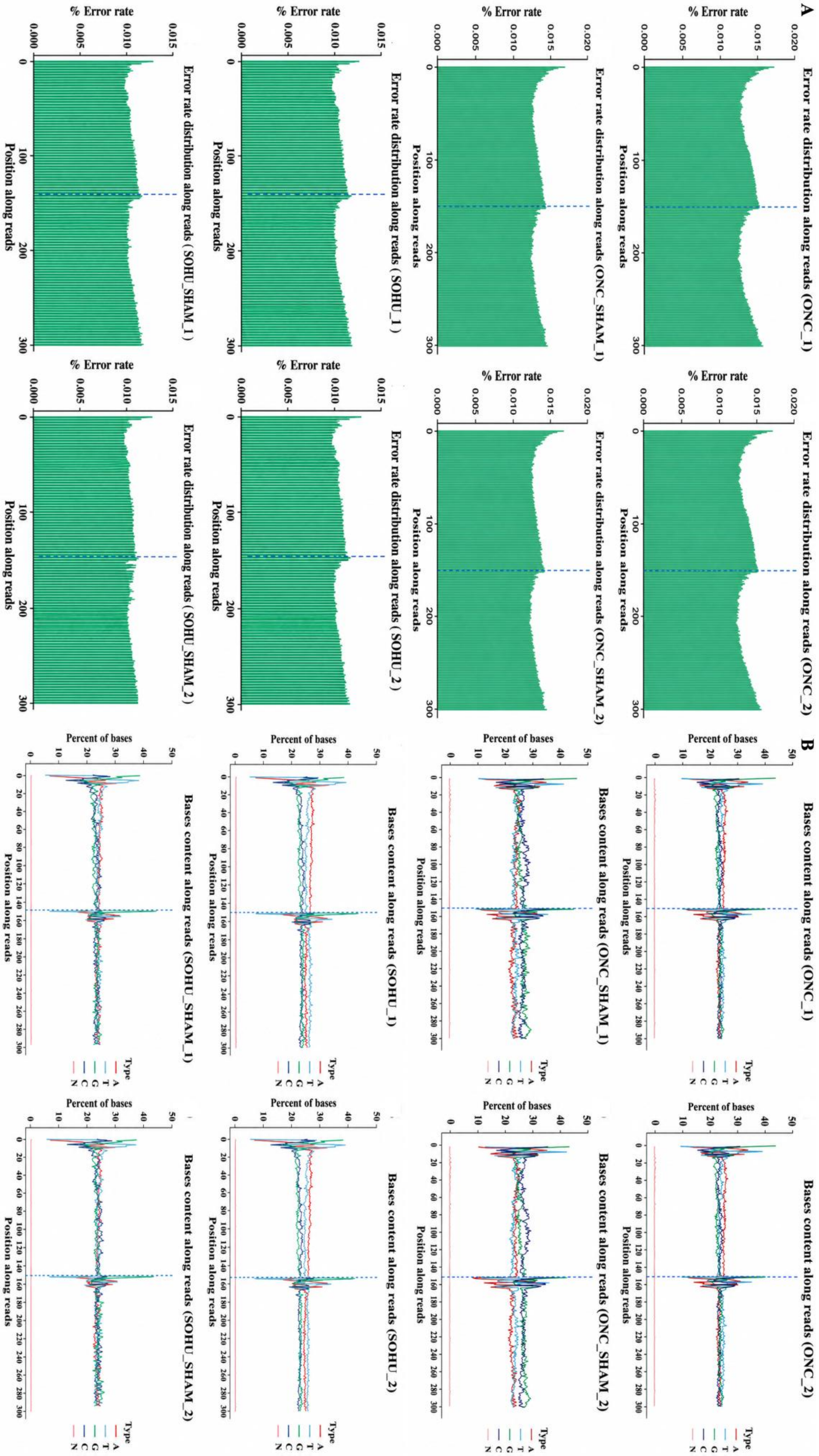


Supplementary Figure 2

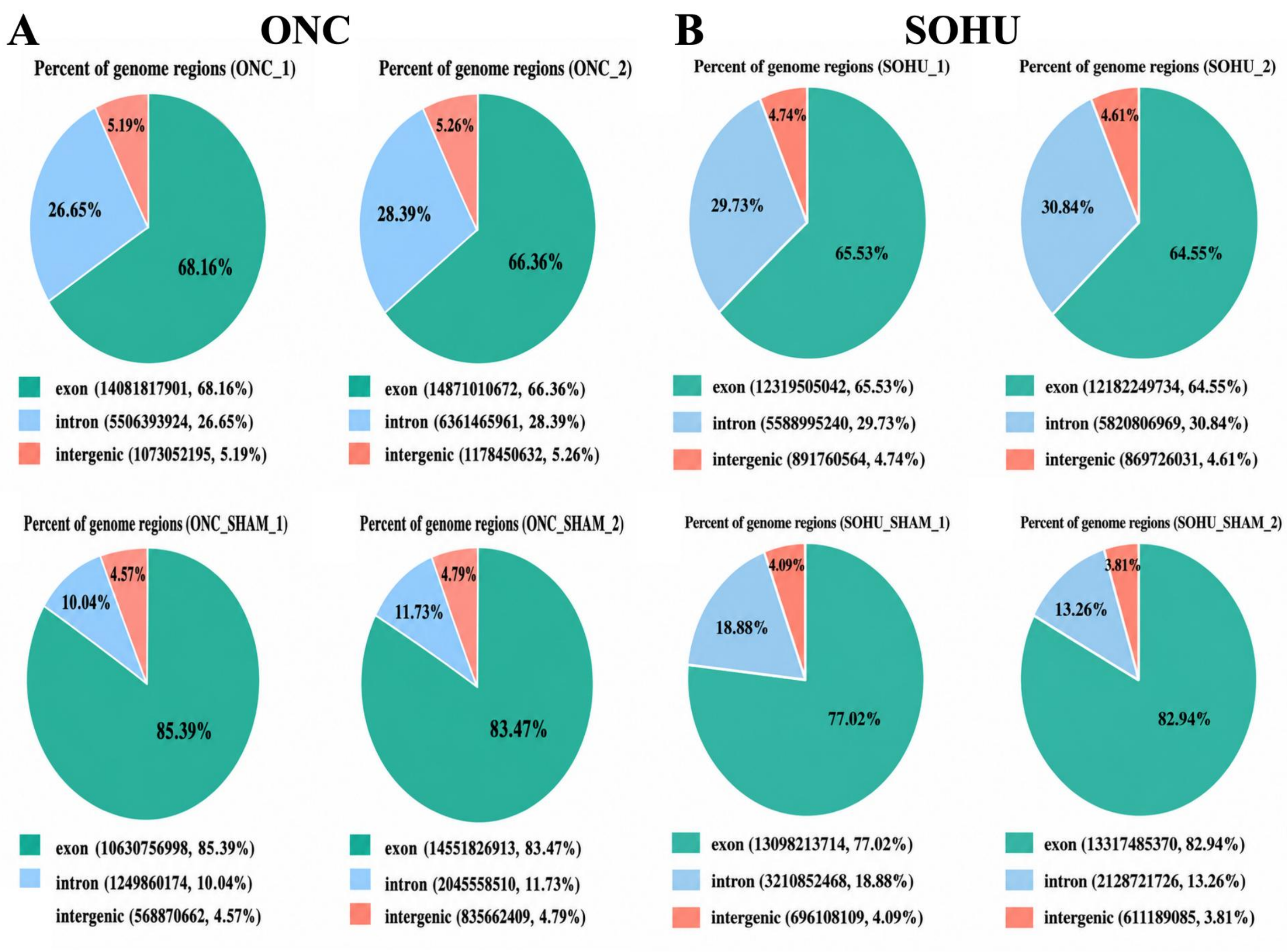


Supplementary Figure 3

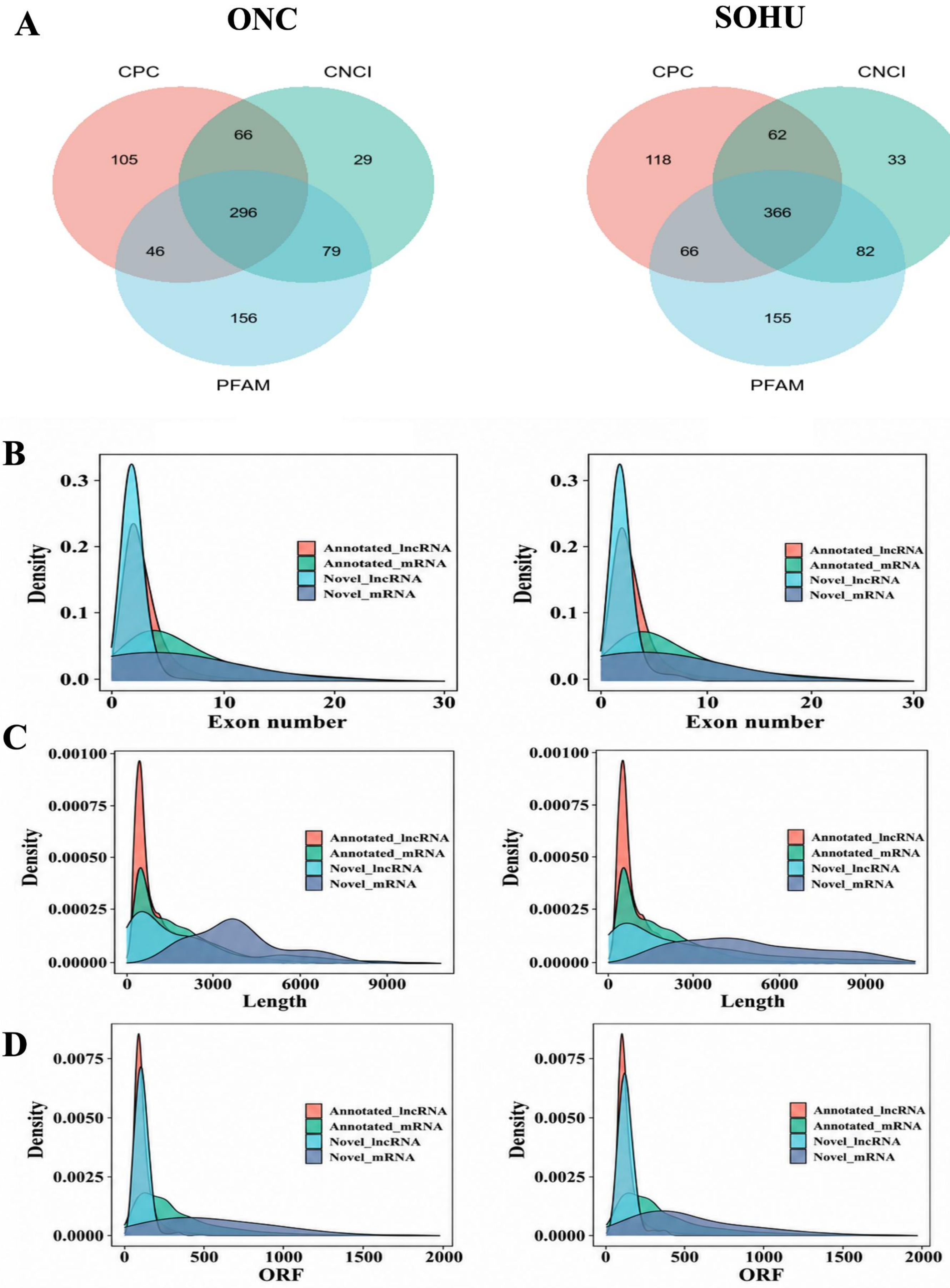


Supplementary Figure 4

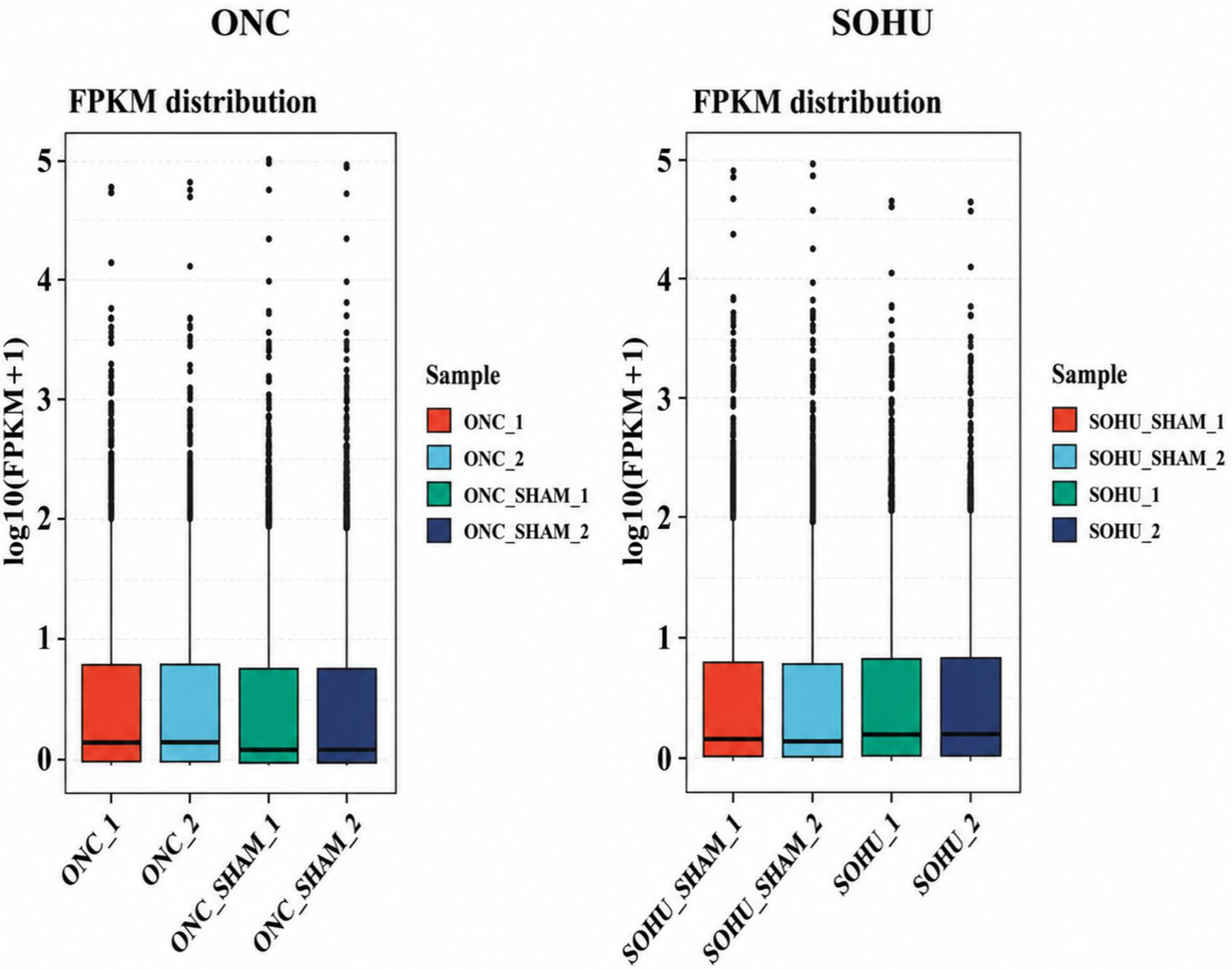


Supplementary Figure 5

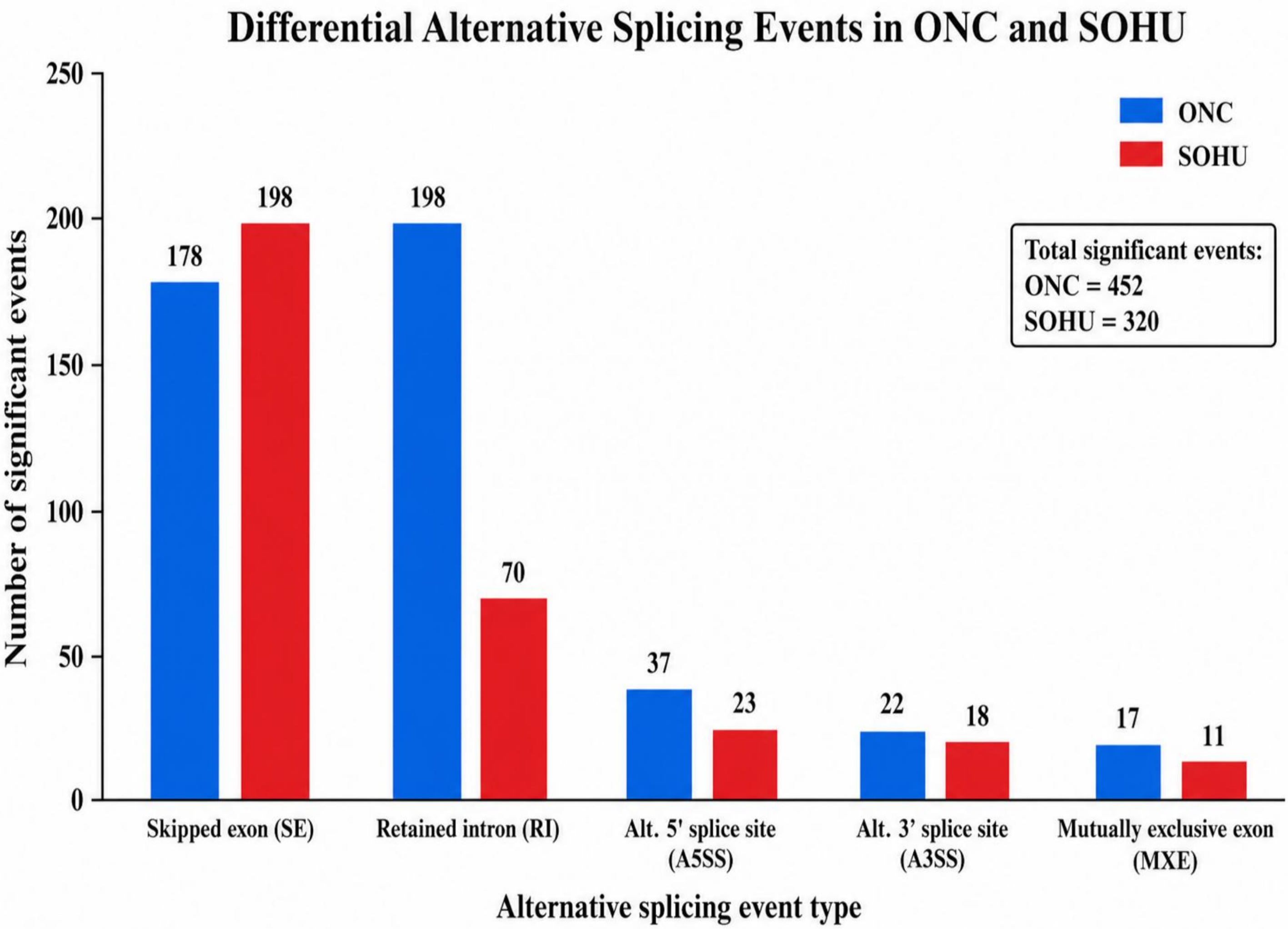


Supplementary Figure 6
